# Single nucleus and *in situ* RNA sequencing reveals cell topographies in the human pancreas

**DOI:** 10.1101/733964

**Authors:** Luca Tosti, Yan Hang, Olivia Debnath, Sebastian Tiesmeyer, Timo Trefzer, Katja Steiger, Foo Wei Ten, Sören Lukassen, Simone Ballke, Anja A. Kühl, Simone Spieckermann, Rita Bottino, Naveed Ishaque, Wilko Weichert, Seung K. Kim, Roland Eils, Christian Conrad

## Abstract

Molecular evidence of cellular heterogeneity in the human exocrine pancreas has not been established, due to the local concentration of hydrolytic enzymes that can rapidly degrade cells and RNA upon resection. Here we innovated single-nucleus RNA sequencing protocols, and profiled more than 120,000 cells from adult and neonatal human donors to create the first comprehensive atlas of human pancreas cells, including epithelial and non-epithelial constituents. Adult and neonatal pancreata shared common features, including the presence of previously undetected acinar subtypes, but also showed marked differences in the composition of the endocrine, endothelial, and immune compartments. Spatial cartography, including cell proximity mapping through *in situ* sequencing, revealed dynamic developmental cell topographies in the endocrine and exocrine pancreas. Our human pancreas cell atlas can be interrogated to understand pancreatic cell biology, and provides a crucial reference set for future comparisons with diseased tissue samples to map the cellular foundations of pancreatic diseases.

## Introduction

Single-cell RNA sequencing (scRNA-seq) has expanded our understanding of heterogeneous human tissues and led to identification of novel functional cell types in the lung, brain and liver (Aizarani et al., 2019; Islam et al., 2014; MacParland et al., 2018; Montoro et al., 2018; Plasschaert et al., 2018). The development of single-nucleus RNA-seq (sNuc-seq) has further broadened application of high-throughput sequencing strategies to tissues which are difficult to dissociate or already archived, including clinical samples (Habib et al., 2017). Pancreatic exocrine tissues contain among the highest levels of hydrolytic enzyme activities in the human body (Farrell, 2010), hindering the preparation of intact RNA from this organ. Therefore, previous scRNA-seq studies of the human pancreas have been restricted to the islets of Langerhans (the endocrine part of the organ) after removal of the exocrine compartment, namely the acinar and ductal cells, the source of digestive enzymes. Following their isolation, endocrine islets were typically cultured *in vitro*, enzymatically dissociated, and processed on microfluidics devices before next-generation sequencing (Baron et al., 2016; Camunas-Soler et al.; Enge et al., 2017; Grün et al., 2016; Lawlor et al., 2017; Muraro et al., 2016; Segerstolpe et al., 2016; Wang et al., 2016). While this strategy proved to be successful in generating a draft of the endocrine human pancreas cell atlas, it has distinct disadvantages. For example, the *in vitro* culture and dissociation steps are known to introduce technical artefacts in gene expression measurements (van den Brink et al., 2017). Moreover, only small numbers of exocrine cells from single cell studies have been reported, leading to underrepresentation of acinar and ductal cells (Muraro et al., 2016; Segerstolpe et al., 2016; Wollny et al., 2016). As a consequence of this underrepresentation, acinar and ductal cells were usually considered homogenous populations dedicated to the production of zymogens and their transport to the intestine, respectively. Thus, the presence, extent or quality of heterogeneity in pancreatic exocrine cells is yet not established.

Here, we innovated methods for rapidly processing tissue biopsies isolated from freshly-isolated human donor pancreata, followed by sNuc-seq, thereby avoiding *in vitro* expansion and dissociation procedures. This approach produced an index draft atlas of human pancreatic cells, including epithelial and non-epithelial cells from both neonatal and adult samples, and revealed previously undetected heterogeneity within pancreatic exocrine cells. Application of *in situ* sequencing combined with computational approaches, enabled us to elucidate spatial relationships and signaling pathways connecting distinct constituent cell types in the pancreas of previously-healthy adult human donors, and revealed dynamic cellular constitution and spatial arrangements during post-natal pancreas development.

## Results

### Innovating sNuc-seq methods for pancreas cells from previously-healthy human donors

To isolate nuclei from frozen tissue, we applied a common protocol based on the use of dense sucrose solutions and detergents at slightly alkaline pH values (Grindberg et al., 2013; Krishnaswami et al., 2016). However, the RNA extracted from isolated nuclei was highly degraded compared to the RNA in the original bulk tissue (Figures S1A-C). Several modifications to the original protocol were systematically applied, including use of dithio-bis(succinimidyl propionate) (DSP) (Attar et al., 2018), methanol fixation (Alles et al., 2017; Chen et al., 2018), or the addition of RNAse inhibitors like ribonucleoside vanadyl complexes (Shieh et al., 2018), but these failed to improve RNA quality. However, on the basis of protocols first described in the 19^th^ century (Carpenter and Smith, 1856) and subsequently modified (Birnie, 1978; Crossmon, 1937; Dounce, 1943), we discovered a citric acid-based buffer which reduced RNA degradation during nuclei isolation, and increased cDNA yields 40-50 fold (compared to standard protocols) from human pancreatic samples (Figure S1D). We isolated nuclei from flash-frozen human pancreas biopsies with short cold-ischemia times (Methods) collected from three male and three female deceased donors, spanning the age range from 1.5 to 77 years (13 samples in total) (Figure 1A and Table S1). The average number of unique molecular identifiers (UMIs) detected per nucleus was 1,287 and the average number of genes detected per nucleus was 692 (Figures S1E). To our knowledge, this effort generated the largest, most comprehensive extant human pancreas cell transcriptome dataset.

**Figure 1.**
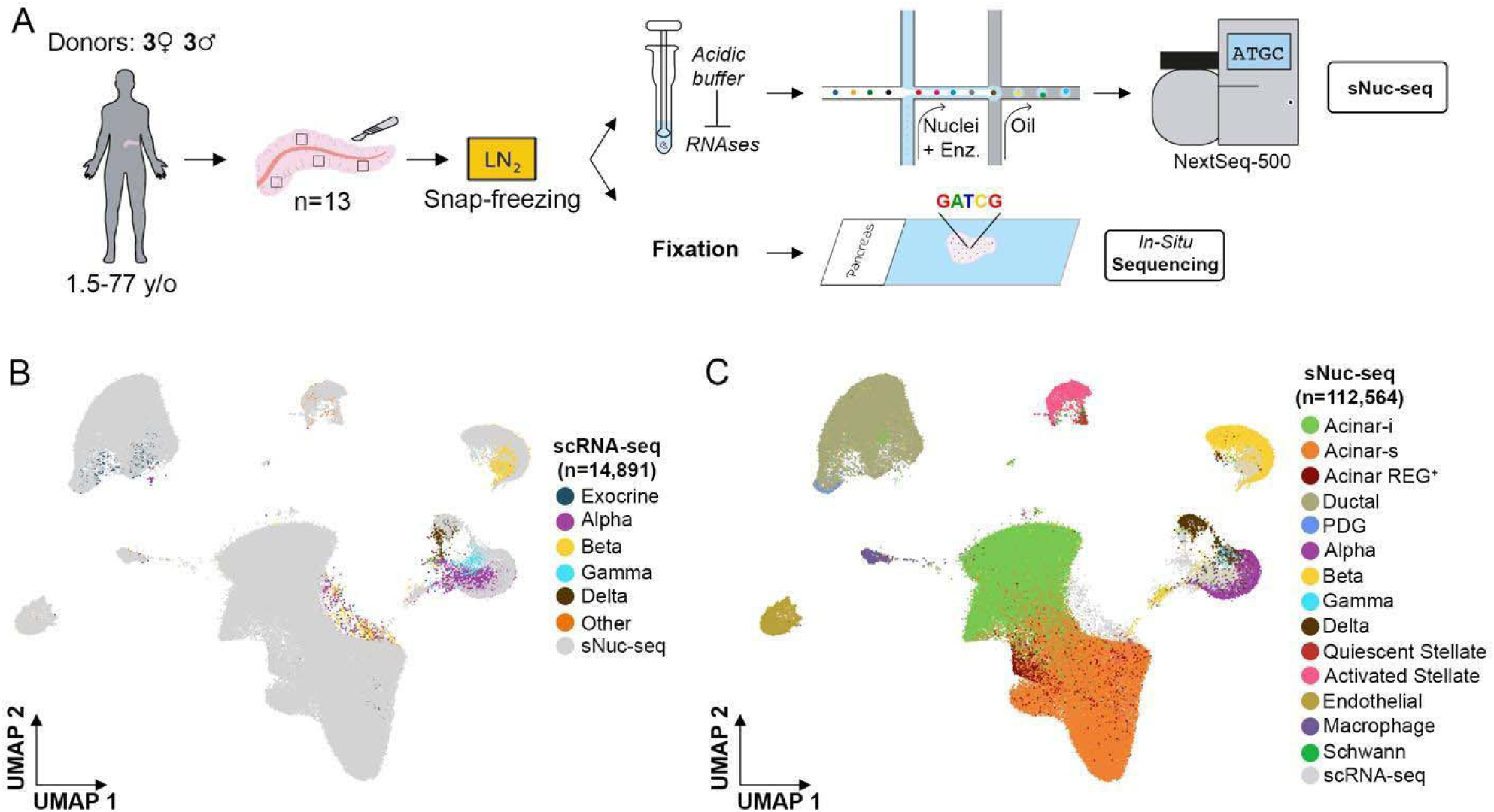
sNuc-Seq identifies cell types in the human healthy pancreas. (A) Overview of the strategy used to perform sNuc-seq and *in situ* sequencing. (B) Merging of sNuc-seq data generated in this study with previous scRNA-seq datasets (Baron et al., 2016; Grün et al., 2016; Lawlor et al., 2017; Muraro et al., 2016; Segerstolpe et al., 2016) of the endocrine human pancreas, shown as clusters in a two-dimensional UMAP embedding. (C) Major cell types identified from sNuc-Seq of the human pancreas shown as clusters in a two-dimensional UMAP embedding.

To aid comprehensive identification and characterization of different constituent pancreatic cell types, we applied canonical correlation analysis (CCA). This achieved (1) reduction of batch effects and (2) integration of data with previously annotated human pancreas scRNA-seq datasets (Figure S1F) (Stuart et al., 2019). Our results confirmed that independent sNuc-seq datasets could be merged and fully integrated with scRNA-seq data sets, despite the use of different starting material (nucleus versus whole cell) (Figure 1B-C) (Mereu et al., 2020). Annotation of cell clusters based on previous studies, confirmed that sNuc-seq enabled us to capture all previously reported pancreatic cell types (Figures 1B and 1C). Moreover, the proportion of cells identified with sNuc-seq differed and complemented data from earlier scRNA-seq studies focused on the endocrine pancreas: in our work, though endocrine cell types were represented, the majority of data derived from acinar or ductal cell nuclei (Figure S2A-B), and also included important non-epithelial cell types (endothelial, stromal, immune cell) not comprehensively characterized in prior work that focused on islet biology.

### Characterization of adult human pancreatic cell types

The two-dimensional UMAP embedding of the sNuc-seq data illustrates distinct cell clusters (Figure 2A). The comprehensive cell representation in our data aligns well with the known composition of the healthy human pancreas, with the majority of analyzed nuclei belonging to two predominant clusters derived from exocrine pancreatic epithelial cells. Acinar cells, accounting for 70% of the nuclei, were identified based on the expression of digestive enzymes such as *CPA1/2, PRSS1* and hallmark transcription factors (TFs) such as *RBPJL* or *FOXP2*. Strikingly, our analysis revealed unanticipated heterogeneity in this cell type (see below). Ductal cells represented 18.5% of the nuclei and expressed cardinal regulators or markers like *CFTR, ANXA4*, and *SLC4A4* (Figure 2B). Unlike in prior studies (Arda et al., 2016; Enge et al., 2017; Muraro et al., 2016; Segerstolpe et al., 2016), we identified two distinct ductal subtypes (Figure 2A) and visual inspection of the principal component loadings confirmed that the two subtypes were separated along the third principal component (Figure 2C). The smaller subtype (accounting for 1% of the total ductal cells) was characterized by higher expression of genes linked to mucous secretion such as the mucin gene *MUC5B* (hereafter, “MUC5B^+^ ductal cells”), the trefoil factor genes *TFF1, TFF2, TFF3*, and the cysteine rich secretory protein 3 *CRISP3* (Figure 2C-D). The other ductal subtype, by contrast, showed higher expression of classical ductal cell markers such as the chloride channel gene *CFTR*, the sodium bicarbonate cotransporter gene *SLC4A4*, and the secretin receptor *SCTR*: collectively, these genes are known regulators of ductal cell secretory function (Figure 2C, 2D) (Baron et al., 2016). Thus, our study provides evidence for unsuspected molecular, and possible functional, heterogeneity in human pancreatic ductal cells.

**Figure 2.**
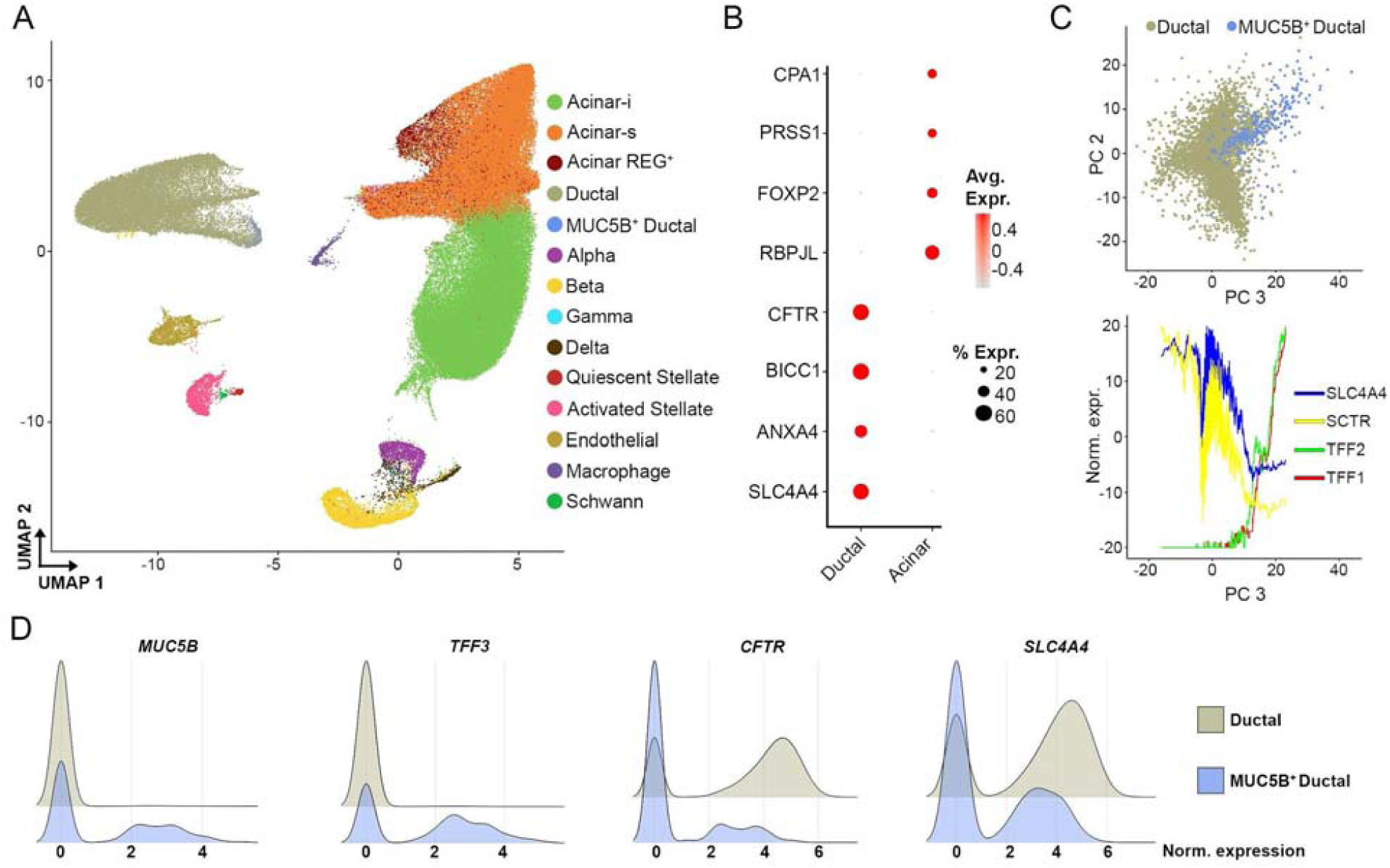
Characterization of ductal cell subtypes. (A) Major cell types identified from sNuc-Seq of the human pancreas shown as clusters in a two-dimensional UMAP embedding. (B) Dotplot showing the expression of specific markers in ductal (including ductal and MUC5B^+^ ductal) and acinar (including acinar-i, acinar-s and acinar-REG^+^) cells. (C) On the top, scatter plot of ductal and MUC5B^+^ ductal cells across the principal component 2 and 3. On the bottom, line plot showing the moving average profile of indicated genes across the principal component 3. (D) Ridge plots showing distinct markers expressed in ductal and MUC5B^+^ ductal cells.

One major group of sNuc-seq clusters contained endocrine cells (approximately 6% of the total number of nuclei) and their identity was confirmed by the expression of known specific hormone genes, namely glucagon (*GCG*, alpha cells), insulin (*INS*, beta cells), pancreatic polypeptide (*PPY*, gamma cells) and somatostatin (SST, delta cells) (Figure 3A). Other clusters included endothelial cells (1.9% of total nuclei), characterized by the expression of *FLT1, PLVAP, VWF, CD36* and *SLCO2A1* and macrophages (0.7% of the total nuclei), expressing *CD74, PTPRC, ZEB2, HLA*-*DRA, HLA-DRB1* and *HLA*-*DPA1* (Figure 3B). We also identified clusters of pancreatic stellate cells (PSCs), which have recognized key roles in normal (Erkan et al., 2012) and diseased pancreas states such as pancreatitis and pancreatic cancer (Shi et al., 2019). We distinguished two distinct states of PSCs in the pancreas from previously-healthy donors, called quiescent (qPSCs) and activated pancreatic stellate cells (aPSCs); based on our cell isolation strategy, these states are unlikely to reflect an artefact of culturing conditions (Baron et al., 2016). qPSCs expressed higher levels of *SPARCL1* mRNA, similar to hepatic stellate cells (Coll et al., 2015), and also *PDGFRB* and *FABP4*, which likely regulate retinoid-storage (Figure 3B and Figure S3) (D’Ambrosio et al., 2011). Moreover, qPSCs were enriched in mRNAs encoding the intermediate filament protein desmin (DES) and integrins such as ITGA1, known regulators of cell structure (Figures 3B and Figure S3). When PSCs activate, they acquire a myofibroblast-like morphology, and are able to migrate and remodel the extracellular matrix (ECM) (Erkan et al., 2012). Both qPSCs and aPSCs express *COL4A1* and *COL4A2*, but aPSCs showed higher levels of mRNAs encoding other collagens such as *COL5A2, COL6A3* and components of the basement membrane such as laminin proteins *LAMA2* and *LAMB1* (Figure 3B and Figure S3). Furthermore, in aPSCs we detected higher mRNA levels of *SLIT2* and *LUM*, known mediators of fibrogenesis and migration in hepatic stellate cells (Bracht et al., 2015; Chang et al., 2015) (Figure 3B and Figure S3). We also detected a cluster of Schwann cells (80 nuclei, 0.02% of total) that expressed characteristic markers like *CDH19, S100B, CRYAB, PMP22* and *SCN7A* (Figure 3B). Over-representation analysis showed the enrichment of specific terms such as “axonogenesis”, “synapse organization” and “synapse assembly” (Figure 3C). By contrast, we did not detect transcripts encoding genes associated with Schwann cell dedifferentiation and reduced myelin sheath formation, that can be upregulated by cell extraction and culture (Baron et al., 2016).

**Figure 3.**
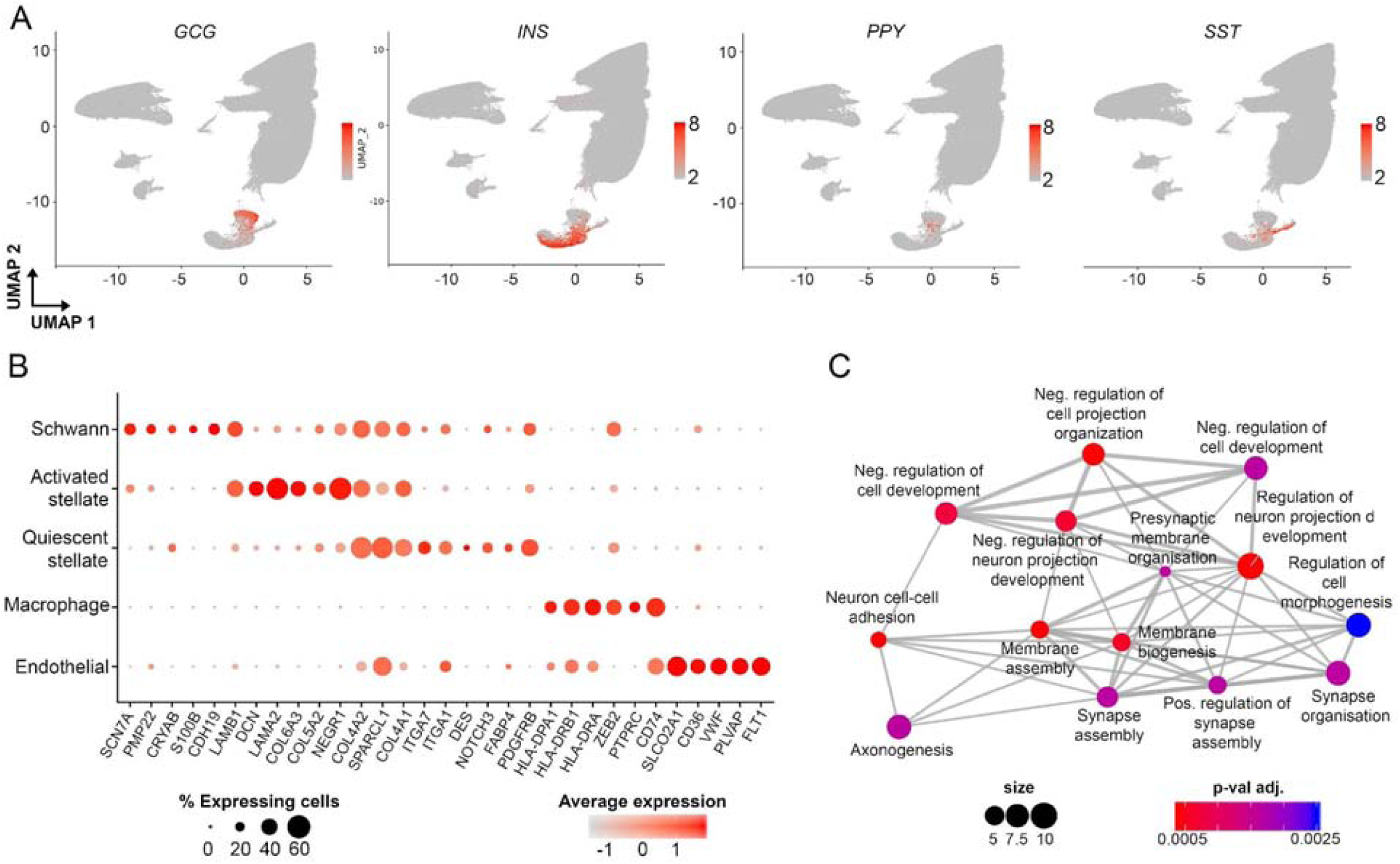
Characterization of other pancreatic cell types. (A) UMAP plots showing the expression of the endocrine cell markers *GCG* (alpha cells), *INS* (beta cells), *PPY* (gamma cells) and *SST* (delta cells). (B) Dotplot showing the expression of specific markers in Schwann, quiescent stellate, activated stellate, endothelial cells and macrophages. (C) Enrichment map of gene ontology terms enriched in Schwann cells.

### Heterogeneity of acinar cells in the adult human pancreas

sNuc-seq data for acinar cells provided an unprecedented opportunity for rigorous assessment of acinar cell heterogeneity, a feature not revealed in prior transcriptomic studies of the human pancreas. One population of acinar cells (acinar-REG^+^) expressed higher levels of mRNAs encoding the regenerating (REG) protein family members such as *REG3A, REG3G* and *REG1B* (Figure 4A and Figure S4). Acinar-REG^+^ cells were reported in a previous scRNA-seq study (Muraro et al., 2016) and represent a population of cells linked to development of pancreatic lesions such as acinar-to-ductal metaplasia (ADM) and pancreatic intraepithelial neoplasia (PanIN) (Li et al., 2016; Liu et al., 2015).

**Figure 4.**
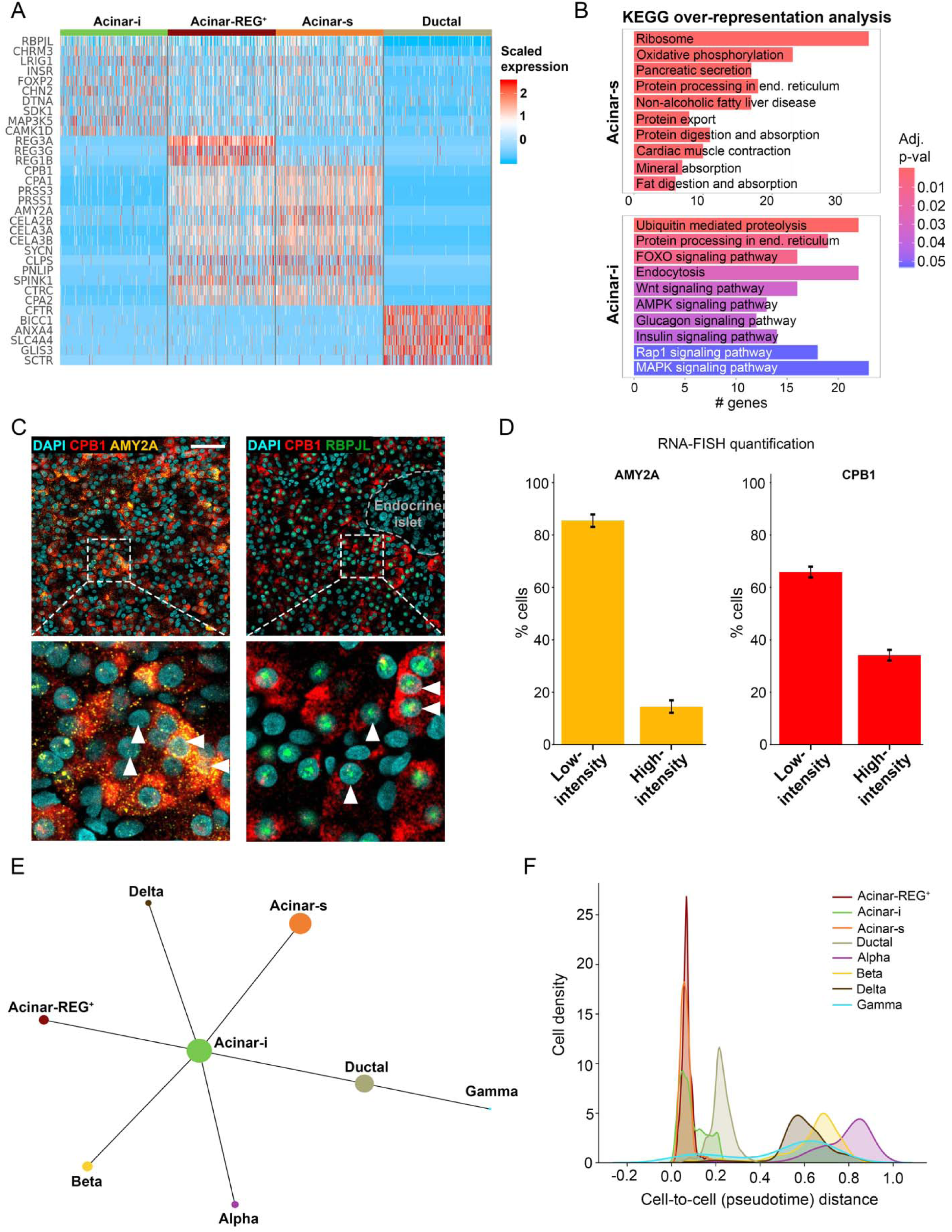
Characterization of acinar cells in the adult human exocrine pancreas. (A) Heatmap of acinar and ductal cell specific genes. (B) Bar plots showing KEGG pathways enriched in acinar-s and acinar-i cells. (C) Example image of RNA-FISH for *CPB1* and *AMY2A*. In the magnified views, the horizontal triangles indicate cells with high intensity RNA-FISH signal, while vertical triangles indicate cells with low-intensity RNA-FISH signal. Scale bar = 50 μm. (D) Quantification of low intensity and high intensity *AMY2A* and *CPB1* RNA-FISH signal in human pancreas sections. The nuclei (n=14,788, 20 images) were classified based on k-means clustering applied to the frequency distributions of pixel counts per nucleus. Error bars indicate standard error of the mean of two independent experiments. (E) PAGA abstracted graph showing the most probable subgraph representing the data manifold. Each node corresponds to a cell type, while the size of nodes is proportional to the number of cells in each cluster. (F) Cell density of pancreatic cell types along a pseudotime trajectory reflecting their transcriptomic similarity.

Strikingly, we detected two additional subtypes of acinar cells not previously identified in human scRNA-seq experiments. These two clusters had distinct UMI levels per nucleus, but a similar number of expressed genes, denoting a distinct complexity of their transcriptomes (Figure S5A). We characterized these two populations by analyzing differentially expressed genes followed by gene over-representation analysis. The acinar cell subtype with higher numbers of UMIs was characterized by higher expression levels of 21 genes (Table S2) encoding for digestive enzymes. Further quantification revealed that 50% of the transcriptome of this cell type encodes for digestive enzyme genes (Figure S5A), confirming previous reports estimating that the majority of the mRNA molecules in a pancreatic acinar cell encode for fewer than 30 proteins (Harding et al., 1977; Hoang et al., 2016). Based on this feature of their transcriptome, we named this subset, “secretory acinar cells” (hereafter, acinar-s). Gene over-representation analysis showed the enrichment of “Ribosome”, “Protein processing in the endoplasmic reticulum” and “Protein export” terms (Figure 4B), consistent with the view that acinar cells have the highest rate of protein synthesis of any human cell (Kubisch and Logsdon, 2008).

The other distinct acinar cell type revealed by our analysis also expressed digestive enzyme genes, but at markedly lower levels compared to acinar-s cells (<4% versus >50%: Figures 4A, S5A). Gene over-representation analysis showed enrichment of terms including “Protein processing in the endoplasmic reticulum”, “Insulin signaling pathway”, “Endocytosis” and “Glucagon signaling pathway” (Figure 4B). Thus, these acinar cells appear less robust in their protein secretion, and instead enriched for responsiveness to external stimuli -like islet signals (Barreto et al., 2010), and activation of the endocytic pathway. We named this subset, “idling acinar cells” (hereafter, acinar-i).

To validate our sNuc-seq findings further, and to evaluate potential role(s) of the acinar-s and acinar-i cells in the healthy pancreas, we combined experimental and computational approaches. First, we performed RNA-FISH on the same samples used for nuclei isolation. Successful RNA-FISH experiments using probes for *CPB1* (Carboxypeptidase) and *AMY2A/B* (Amylase) were performed and after quantification (Figure S5C) we confirmed the existence of distinct acinar cells expressing different levels of digestive enzyme genes (Figure 4C-D). In particular, mRNA of *CPB1* and *AMY2A/B* showed heterogeneity across the tissue and, in agreement with sNuc-seq results, we were able to distinguish two classes characterized by differential RNA-FISH signal (Figure 4C-D).

Second, we applied SCENIC, a computational tool for inferring transcription factor-target regulatory networks (regulons) from single cell gene expression (Aibar et al., 2017). Both the acinar-s and acinar-i subtypes showed activation of the regulon *CREB3L1*, likely involved in the basal secretory activity of the cells (Figure S5B). The *XBP1* regulon shows high activation in acinar-s cells in agreement with the role of this transcription factor in the unfolded protein response (UPR) pathway (Lee et al., 2003), and consistent with the view that biosynthetic activity and accompanying increase of endoplasmic reticulum (ER) stress are higher in acinar-s cells (Figure S5B). Notably, only acinar-s cells showed activation of regulons associated with the maintenance of acinar cell identity such as *GATA4, NR5A2* and *MECOM* (Figure S5B).

Third, we elucidated relationships between acinar-s or acinar-i subtypes and other pancreatic cell types using partition-based graph abstraction (PAGA) (Wolf et al., 2019). With PAGA representation, nodes represent distinct cell “states”, while edges indicate potential routes of cell transitions between them. Here, we included cell types which are known to derive from a common multipotent progenitor during embryonic development (Zhou et al., 2007), namely acinar and ductal cells in the exocrine compartment, and alpha, beta, gamma and delta cells in the endocrine compartment. This unsupervised approach places acinar-i cells in a *central* position, showing similar connections to the majority of the other cell types such as ductal and endocrine cells (Figure 4E). Fourth, we exploited the principle of pseudotime analysis to order cells on the basis of the similarity of their transcriptome (Trapnell, 2015). Interestingly, acinar-i cells occupy an intermediate position between acinar and ductal cells, reflecting the known plasticity of acinar cells and their ability to convert towards the ductal lineage (Figure 4F) (Storz, 2017).

### Single nucleus sequencing of the human neonatal pancreas

A wealth of data is available about the embryonic development of pancreas in mammals (Jennings et al., 2015; Kim et al., 2020; Larsen and Grapin-Botton, 2017), but much less is known about the postnatal development of this organ. Here, we procured two 1-day old samples (Table S3) from one male and one female donor and generated sNuc-seq data from 10,528 nuclei, with an average number of UMIs and genes per nucleus of 964 and 628, respectively (Figure 5A). We identified different cell types and cellular compositions specific for this developmental stage (Figure 5B-C). In particular, the exocrine neonatal compartment (acinar and ductal cells) accounts for around 50% of the organ, while it constitutes about 90% of the adult pancreas, in agreement with early studies of postnatal growth performed in rodents (Figure 5C) (Elsässer et al., 1994; Kachar et al., 1979; Sidorova and Babaeva, 1968). Moreover, at the postnatal stage, pancreatic endocrine cells account for 22% of detected cells in neonates compared to 6% in the adult (Figure 5C). Endocrine cells showed changes in cellular composition, with neonatal delta cells, the third major cell type of endocrine islets, accounting for 29% of the endocrine cells compared to 15% in adults (Figure 5D) (Rahier et al., 1981; Stefan et al., 1983). Alpha and beta cells accounted for 21% and 48% of the neonatal endocrine cells, in line with findings in humans and pigs (Figure 5D) (Kim et al 2020; M. Brissova, A. Powers, S.K. Kim, unpubl. results). While we did not detect a gamma cell population, we captured 42 *GHRL*^*+*^ (Ghrelin) epsilon cells in the endocrine compartment (Figure 5B) (Wierup et al., 2002). Ghrelin binds to cell-surface receptors like GHSR, and is known to play an insulinostatic function (DiGruccio et al., 2016; Reimer et al., 2003). Differential gene expression analysis of epsilon cells revealed that after ghrelin (*GHRL*), the second most differentially expressed gene is *ACSL1*, encoding for an Acyl-CoA synthetase enzyme which catalyzes the unique addition of an octanoyl group to ghrelin, a modification which is essential for its optimal biological activity (Gutierrez et al., 2008; Hougland, 2019). Other epsilon cell markers identified in this study include the annexin family member *ANXA13*, the proline-, histidine-, the calcium permeant cation channel *TRPC4* and the asialoglycoprotein receptor *ASGR1* (Figure 5E).

**Figure 5.**
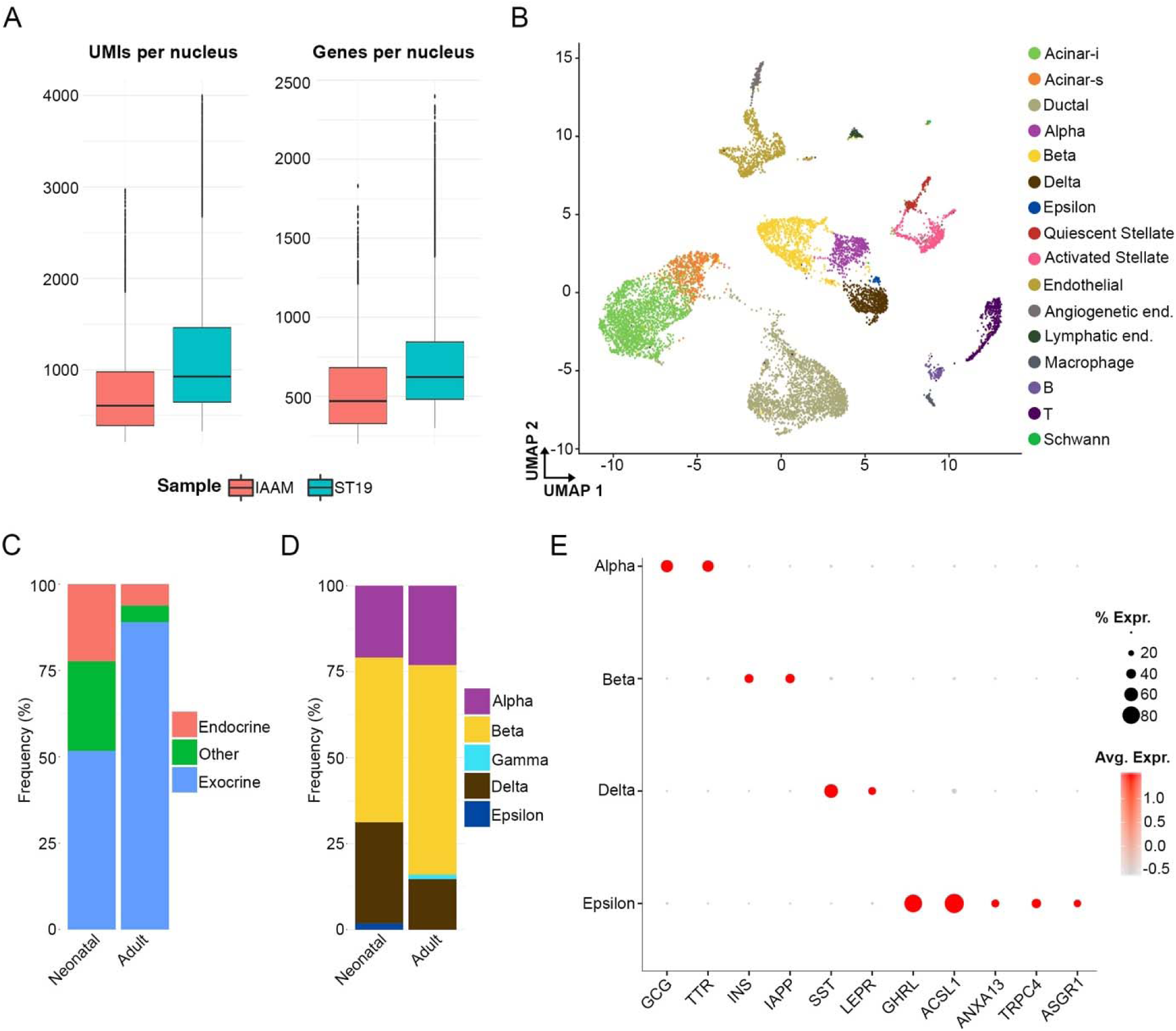
Characterization of the cellular composition of neonatal healthy pancreas. (A) On the left, the boxplots show the distribution of UMIs per nucleus for each neonatal sample processed in this study. On the right, the boxplots show the distribution of genes per nucleus for each neonatal sample. (B) Major cell types identified from sNuc-Seq of the human neonatal pancreas shown as clusters in a two-dimensional UMAP embedding. (C) Frequency of different cell types in adult and neonatal pancreas. (D) Frequency of different endocrine cells in the adult and neonatal pancreas. (E) Dot plot of distinct genes expressed in neonatal endocrine cells.

In exocrine cells from neonates, like in adults, we detected acinar-i and acinar-s cells, including similarities of UMI and gene count distributions (Figure 6A). For example, 4% of the transcripts of neonatal acinar-i cells encode for digestive enzyme genes compared to 33% in acinar-s, similar to the levels found in cognate adult cells (Figure 6A). However, we did not detect *AMY2A, AMY2B* and *PNLIP* in neonatal acinar-s cells, consistent with prior findings that little to no pancreatic amylase or lipase enzyme activity is detectable in newborns (Figure 6B) (Lebenthal and Lee, 1980; Zoppi et al., 1972). Moreover, we did not detect a population of acinar-REG^+^ cells in neonatal samples, suggesting a further layer of REG protein regulation during postnatal to adult maturation (Figure 5B). In the ductal compartment, we did not detect the MUC5B^+^ population, but this may reflect low duct cell yields from human neonatal pancreas (Figure 5B).

**Figure 6.**
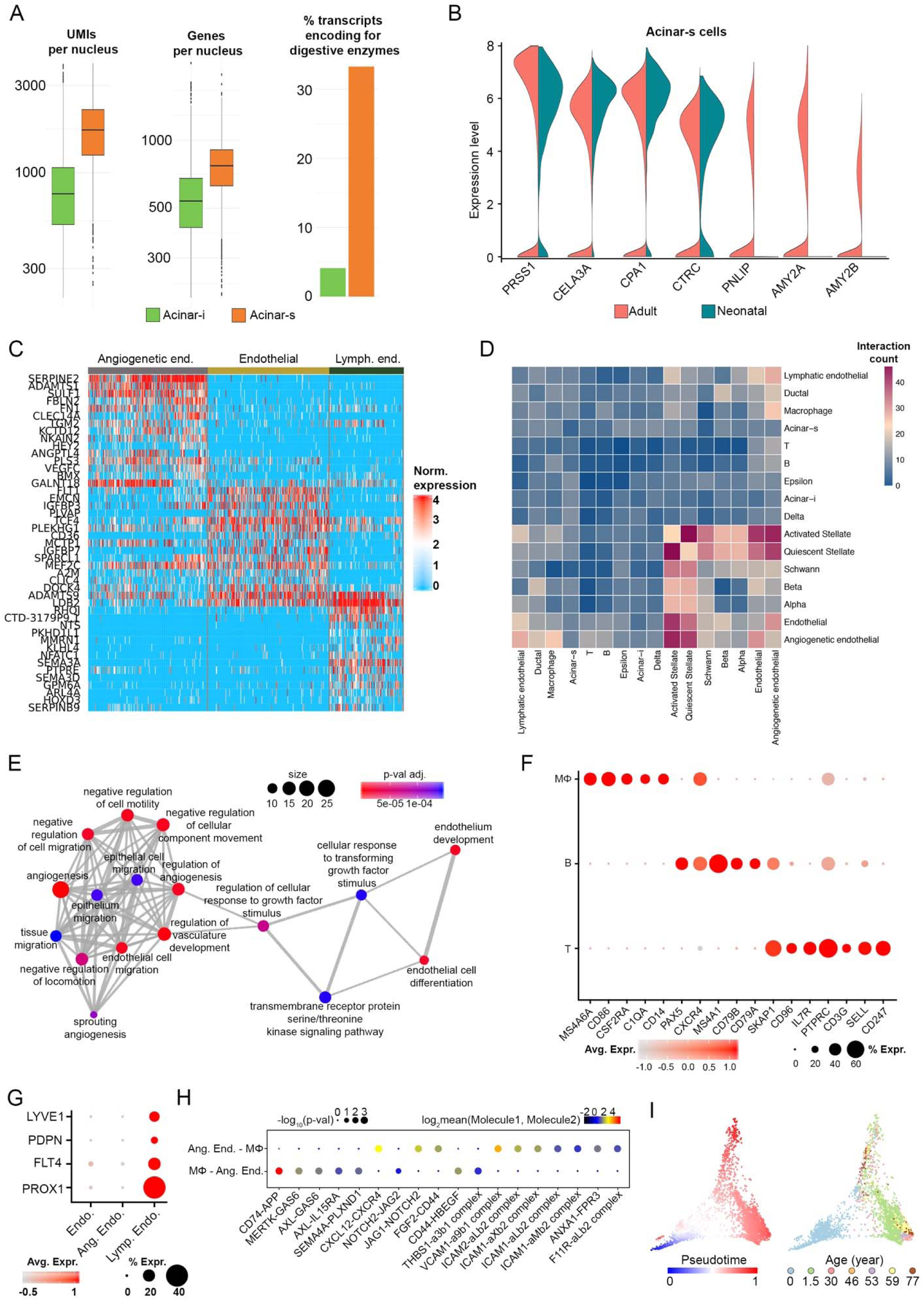
Characterization of the acinar, endothelial and immune constituents of the neonatal healthy pancreas. (A) Quantification of UMI per nucleus (left) and genes per nucleus (center) for the different acinar cell states. On the right, the percentage of transcriptome encoding for digestive enzymes (Table S2) is represented. (B) Violin plot showing the expression level of selected digestive enzyme genes in adult and neonatal pancreas. (C) Heat map showing different genes expressed in three different endothelial cell types. (D) Heat map depicting the number of all possible interactions between the analyzed cell types of the neonatal pancreas as calculated by CellPhoneDB. (E) Enrichment map of gene ontology terms enriched in angiogenetic endothelial cells. (B) Dot plot of specific lymphatic endothelial cell markers. (F) Dot plot showing the expression of specific genes in macrophages (MΦ), B and T cells. (G) Dot plot of specific lymphatic endothelial cell markers. (H) Dot plot depicting selected MΦ-angiogenetic endothelial interactions enriched in healthy neonatal pancreas. (I) Scatter plot in Diffusion map basis (components 1 and 2) of combined neonatal and adult beta cells, colored by pseudotime (left) or by age of the donor (right).

Unlike in the adult pancreas, we observed evidence of endothelial cell heterogeneity in neonatal samples (Figure 6C). For example, in addition to an “adult-like” endothelial signature, we observed an “angiogenetic” endothelial type enriched for mRNA previously linked to programs of blood vessel morphogenesis and extracellular matrix remodeling (Figure 6E). Furthermore, on the basis of the expression of canonical markers such as *LYVE1, PDPN, FLT4* and *PROX1*, we also identified a population of *lymphatic* endothelial cells that was not detected in the adult pancreas (Figure 6G) and that most likely functions to collect interstitial fluid containing cell debris (Cesmebasi et al., 2015; O’Morchoe, 1997).

Few immune cells were detected in the human adult healthy pancreas, even though localization of immune cells to the pancreas can be dramatically altered in pancreatic diseases (Zheng et al., 2013). By contrast, in the neonatal pancreas we identified an abundance of at least three immune populations including follicular B cells (*CD19, CR2, CD22, FCER2*), T lymphocytes (*CD247/CD3Z, CD3G, IL7R*) and macrophages (*CD14, CD86, CSF2RA*) (Figure 5B and 6F). To further clarify the potential cell-cell interactions of macrophages with other cell types in the neonatal pancreas, we applied CellPhoneDB, a statistical framework used to predict cellular interactions (ligand-receptor) from single-cell transcriptomics data (Vento-Tormo et al., 2018). Analysis of cell-cell interactions revealed that macrophages have a higher number of interactions with angiogenic endothelial cells (Figure 6D), supporting the view from studies of other organs that macrophages regulate vascular development and remodeling (Fantin et al., 2010; Nucera et al., 2011). Among the ligand-receptor interactions revealed by CellPhoneDB between macrophages and angiogenetic endothelial cells, we noted specific sets of ligands and receptors, such as the TAM receptors (*AXL, MERKT*, and the ligand *GAS6*), usually active in tissues subject to remodeling and involved in the phagocytosis of apoptotic cells (Lemke, 2013) and members of the Notch pathway including the *NOTCH2* receptor and the antagonistic ligands *DLL4, JAG1* and *JAG2* (Pitulescu et al., 2017). Moreover, a strong interaction was predicted between *CD74* and *APP*, suggesting an important role for *APP* in pancreatic neonatal angiogenesis (Figure 6H).

We next used sNuc-seq data to investigate age-dependent development, with a specific focus on beta cells. Beta cells are not fully functional at the perinatal stage but their functions, including glucose-regulated insulin secretion, mature with age (Arda et al., 2016). To investigate changes across the two different age groups (neonatal and adult), we combined the two datasets and performed diffusion pseudotime analyses (Haghverdi et al., 2016). Pseudotime ordering recapitulated donor age (Figure 6I), thereby permitting us to model gene expression using a generalized additive model and to identify highly dynamic genes. We identified groups of genes showing increasing or decreasing expression levels across pseudotime (Figure S6A). Some of the genes expressed at lower levels in adult samples are involved in beta cell proliferation such as *PDZD2, IGFBP5* and *CDK6* (Blum et al., 2012; Gleason et al., 2010; Ma et al., 2006; Suen et al., 2008; Takane et al., 2012) (Figure S6A-B). PLAG1, a protein known to decline within a few days after birth and known to inhibit insulin secretion in neonatal murine islets (Hoffmann and Spengler, 2012), was also detected in neonatal samples (Figure S6A-B). By contrast, *CD99*, whose expression increases in adult mouse islets (Aguayo-Mazzucato et al., 2017), also had increased mRNA levels in adult human beta cells (Figure S6A-B). In human adults, we also detected higher levels of *RASD1* and *SYT16*, genes previously reported to be upregulated in human islets exposed to relatively high glucose (Hall et al., 2018; Huang et al., 2018). Genes encoding members of the secretogranin-chromogranin family like *SCG2, SCG5* and *CHGB*, known for their essential role within the insulin secretory granule, were also more highly expressed in adult beta cells (Bearrows et al., 2019; Obermüller et al., 2010; Suckale and Solimena, 2010) (Figure S6A-B). Thus, by applying sNuc-seq to tissues procured at specific developmental stages – our work reveals markers and unrecognized possible regulators of human beta cell functional maturation.

### *In situ* sequencing of the human pancreas localizes mRNA

To elucidate the role of the different heterogeneous cell states identified in our study we integrated our sNuc-Seq results with *in situ* sequencing (ISS) using matching tissues processed for nuclei isolation (Figure 1A). ISS combines the use of padlock probes and rolling circle amplification directly in tissue sections, permitting targeted measures of expression by genes of interest, and identification of cell markers and cell subtypes at single cell resolution (Figure S7A) (Ke et al., 2013; Qian et al., 2019). In our analysis we selected 83 marker genes (Table S4) identified from sNuc-seq that distinguish pancreatic cell types, then applied ISS to tissue from one juvenile (1.5 years old donor) and two adult (30 and 53 years old) donors. ISS-based mRNA localization (Figure S7A) was used to generate spatial cell maps by applying Spot-based Spatial cell-type Analysis by Multidimensional mRNA density estimation (SSAM), a segmentation-free algorithm that identifies cell-type signatures from spatially resolved *in situ* transcriptomics data (Figure 7A and Figure S7B) (Park et al., 2019).

**Figure 7.**
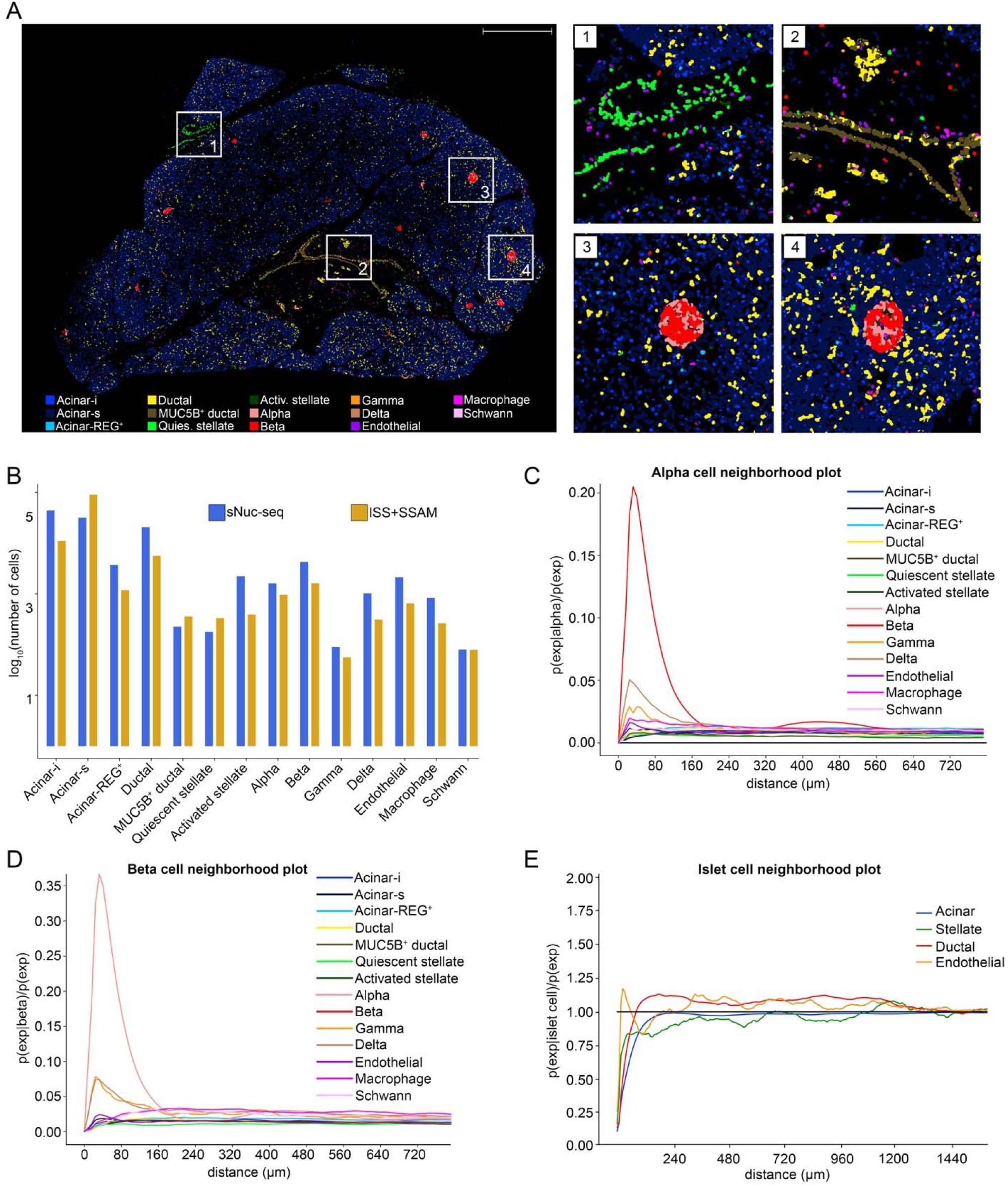
*In situ* sequencing of the human healthy pancreas. (A) On the left, the cell map generated by SSAM from a tissue section of an adult donor (AFHE365-head). On the right, magnified views showing macroscopic features such as (1) quiescent and activated stellate cells in the connective tissue, (2) interlobular duct enriched in MUC5B^+^ ductal cells, (3) and (4) endocrine islets enriched in alpha and beta cells, but also including delta, endothelial cells and macrophages. Scale bar = 1 mm. (B) Bar plot comparing the numbers of cells identified via sNuc-seq with the normalized surface area calculated by SSAM for each cell type. (C) Line plot showing the results of the spatial modelling analysis for the alpha cells. (D) Line plot showing the results of the spatial modelling analysis for the beta cells. (E) Line plot showing the results of the spatial modelling of islet cells (combination of alpha, beta, gamma and delta cells). Stellate include “activated” and “quiescent” stellate cells, ductal includes “ductal” and “MUC5B^+^ ductal” cells, acinar includes acinar-i, acinar-s and acinar-REG^+^ cells.

SSAM cell maps contained all the cell types identified by sNuc-seq, and permitted ready recognition of multicellular tissue features including endocrine islets, interlobular ducts and stellate cells in the connective tissue of septae (Figure 7A, magnified views). The proportion of cell types detected by SSAM cell maps also corresponded well with those detected using sNuc-seq (Figure 7B), further confirming the robustness of our sNuc-Seq analyses. To probe spatial relationships between pancreatic cell types, we performed empirical statistical modelling of cell type proximity (Methods). Initially, to confirm the validity of this approach, we quantified spatial relations within unambiguous multicellular structures. For example, SSAM modelling results confirmed the mutual proximity of alpha and beta cells (Figure 7C-D), reflecting their characteristic localization within the islets. Furthermore, the delta and gamma cells (also part of the endocrine islets, but in much smaller numbers) are the second and third most likely cell types to be found within a distance of 20-80 μm from alpha and beta cells (Figure 7C-D). We looked at non-epithelial cell types in close proximity to the endocrine islets and found that endothelial cells are the closest ones (Figure 7E), as they support high oxygen demand, and the glucose-sensing and endocrine functions of islets (Bonner-Weir and Orci, 1982; In’t Veld and Marichal, 2010). Analysis of ISS provided further insights into the intra- and inter-islet architecture. Quantification of islet size revealed a higher frequency of small islets (radius smaller than 40 μm) in juvenile tissue compared to the adult (Figure 8A). We then performed proximity analysis on cells located outside of the islets and discovered that, in the juvenile sample (1.5 years), alpha cells are the most proximal (in the first 40 μm) followed by beta cells, suggesting enrichment of alpha cells in the mantle of the islets, while beta cells preferentially locate in the core (Figure 8B) (Bonner-Weir et al., 2015; Bosco et al., 2010). Importantly, this trend is diminished in adult samples (30 and 53 years), reflecting an age-dependent increase of architectural heterogeneity in adult islets (Figure 8B) (Dybala and Hara, 2019). We then calculated the distance between the centroids of manually annotated endocrine islets; in both neonatal and adult samples, we found a minimum distance of 400 μm, providing quantification of pancreatic islet dispersion during pancreas morphogenesis (Figure 8D) (Hastings et al., 1992; Pauerstein et al., 2017).

**Figure 8.**
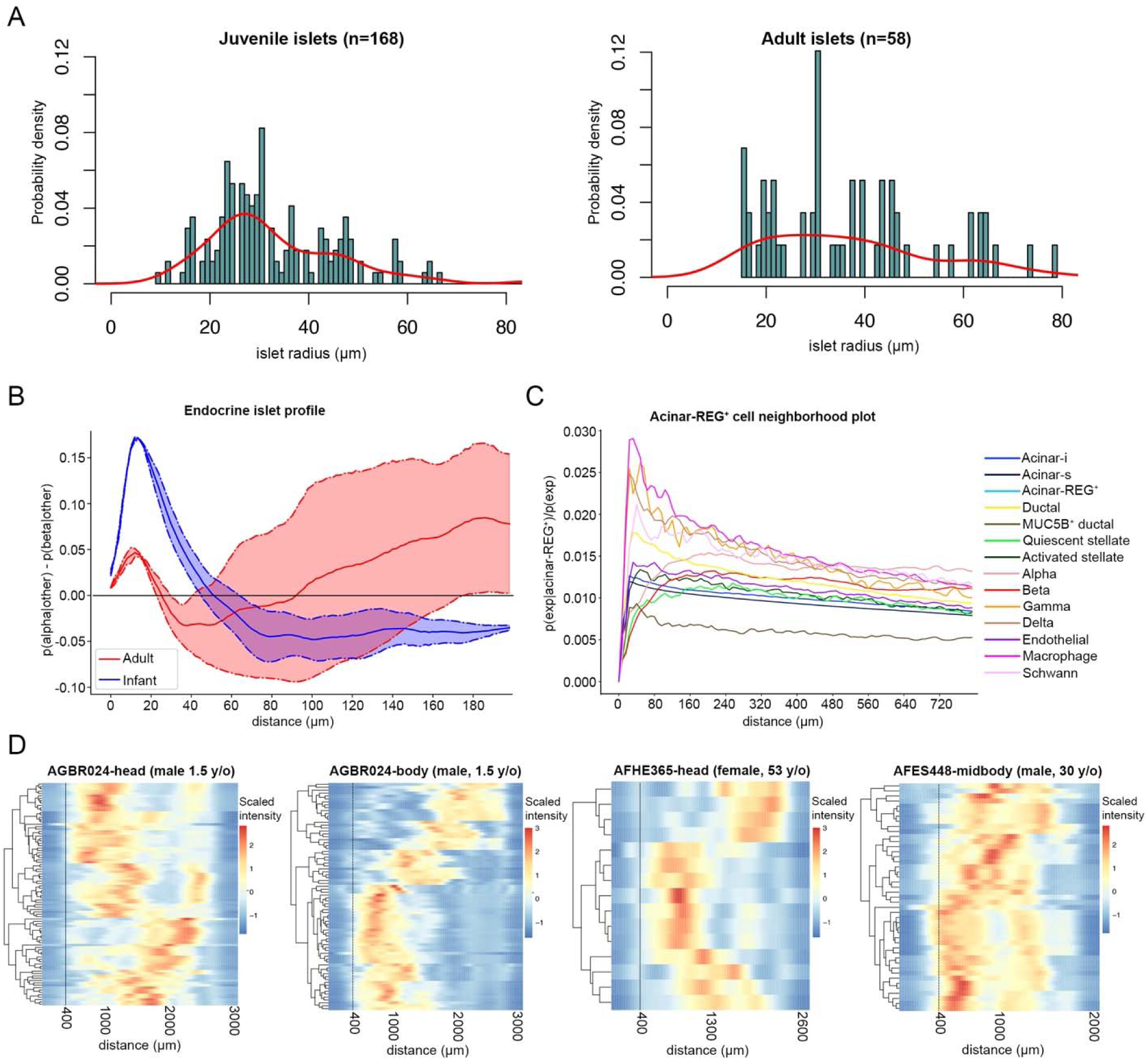
Characterization of intra and inter-islet architecture. (A) On the left, histogram and density line showing the distribution of juvenile islet radii. On the right, histogram and density line showing the distribution of adult islet radii. (B) Line plot showing the results of the spatial modelling analysis for any cell surrounding the endocrine islets. (C) Line plot showing the results of the spatial modelling analysis for the acinar-REG^+^ cells. (D) Each row of the heatmaps represent a single islet in each sample. The distances between the centroids of each islet and all the other islets were calculated and the scaled intensity of the frequency is represented in each row. High (red) and low (blue) values indicate higher or lower presence of other islets at the specific distance, respectively.

We also investigated the spatial relations of the new acinar cell states identified in this work. In particular, the acinar-s and the acinar-i cells did not show a specific cell neighborhood (Figure S8), a result in agreement with their vast abundance (about 80%) in the tissue. By contrast, acinar-REG^+^ cells appeared to localize significantly closer to islet cells, like delta and gamma cells (Figure 8C) (Muraro et al., 2016), and to macrophages. This latter finding is consistent with prior reports that REG3A/PAP protein modulates chemoattraction and activation of macrophages in pancreatic disease and in neural tissues (Gironella et al., 2013; Namikawa et al., 2006; Viterbo et al., 2008). Together, these results highlight how mRNA localization combined with sNuc-seq can be used to identify cell types and reconstruct known and unrecognized morphological patterns in the pancreas.

## Discussion

Here, we constructed a comprehensive human pancreas cell atlas by combining high-throughput nuclear RNA sequencing and RNA localization. To achieve this, we developed novel strategies for nucleus isolation from human pancreas that could be applied to other challenging tissues and to archived clinical samples. Moreover, we successfully generated the first *in situ* sequencing dataset of the juvenile and adult pancreas. These approaches revealed unsuspected heterogeneity in pancreatic cells and cellular interactions, including age-dependent cellular arrangements. Together, our findings provide an unprecedented, comprehensive resource for the community of science focused on pancreas and organ biology.

The heterogeneity of the exocrine pancreas has been previously investigated using immuno histology-based assays (Adelson and Miller, 1989; Uchida et al., 1986), but recent single cell analyses (Muraro et al., 2016; Segerstolpe et al., 2016; Tritschler et al., 2017; Wollny et al., 2016) have suggested that high-throughput sequencing could be a powerful tool to reveal undetected singularities in pancreatic exocrine cell types. However, the inference of pancreatic exocrine cell heterogeneity in prior work was based on relatively small sample sizes, reflecting the primary focus of these studies on islet biology (Wollny et al 2016: Baron et al 2016). From studies of over 120,000 pancreatic cells, we found evidence of three distinct acinar cell populations (acinar-i, acinar-s and acinar-REG^+^), distinguished by differential expression of digestive enzyme genes, distinct activation of pancreatic gene regulatory networks, expression of specific protein family genes and distinct cell neighborhoods. *In vitro* systems (like cell lines or organoids) that reconstitute mature human acinar cells in their native architecture have not yet been achieved, therefore precluding functional validation analyses in such systems. Instead, we used orthogonal approaches to validate our sNuc-seq results and to infer cellular and signaling mechanisms, including a combination in human pancreas of RNA-FISH, *in situ* sequencing and modern computational approaches. RNA-FISH and *in situ* sequencing approaches have not been robustly applied to human pancreas since, as for sNuc-seq, elevated RNA degradation typically hinders these experiments. Together, the assays performed in this study elucidated the potential role of the distinct acinar cell states we identified.

Our studies revealed that acinar-REG^+^ cells were absent in the neonatal pancreas, suggesting that their function might be specific for the adult tissue. Previous scRNA-seq studies described a subset of acinar cells with lower expression levels of digestive enzyme genes and their localization around the endocrine islets (Muraro et al., 2016). Those findings were recapitulated by our sNuc-seq. Moreover, application of *in situ* sequencing analyses - for the first time, to our knowledge - in the human pancreas, revealed significant localization of acinar-REG^+^ near macrophages, nominating acinar-REG^+^ cells as possible regulators of pancreatic inflammatory processes. The acinar-s cell state conforms to a “classical” view of the pancreatic acinar cell, which is characterized by a specific gene regulatory network and mainly committed to the production, processing and regulated secretion of pancreatic zymogens. In acinar-i cells we find evidence suggesting that hydrolytic enzyme production may be reduced, compared to acinar-s cells. We speculate that this acinar cell “state” (Morris, 2019) could be a protective adaptation to periods of intense zymogen production and increased endoplasmic reticulum stress. If so, acinar-i cells might be analogous to a subset of postulated metabolically-stressed islet beta cells (Baron et al., 2016; Szabat et al., 2016; Xin et al., 2018). Acinar-i cells showed a decreased activation of acinar cell gene regulatory networks and occupied a central position in a PAGA lineage relation graph, and we speculate that these cells might have the capacity to convert into other pancreatic cell types, including both ductal and endocrine cells (Stanger and Hebrok, 2013). Further investigations should clarify how acinar heterogeneity is achieved and maintained during homeostasis, whether the acinar-i, acinar-s and acinar-REG^+^ cells can interconvert under physiological conditions, and what - if any - impact the acinar cell heterogeneity has in development of pancreatic exocrine disorders like pancreatitis, acinar-to-ductal metaplasia, and pancreatic ductal adenocarcinoma. Here, we did not identify centro-acinar cells, which have been postulated to include pancreatic progenitor cells (Rovira et al., 2010). Future studies could clarify the transcriptome of centro-acinar cells in the human pancreas, and their possible lineage or spatial relation with the acinar cell and ductal cell types captured in our datasets.

Here we also demonstrated the feasibility of deploying our nuclei isolation strategy for pancreas from children and neonates. This has revealed developmental dynamics in the pancreas, unique immune interactions (involving B, T cells, and macrophages), and changes in cellular composition compared to the adult pancreas. For example, within the endocrine compartment we detected a higher number of delta cells compared to those in the adult. Since somatostatin output by delta cells is known to inhibit cell proliferation and to promote apoptosis (Patel and Srikant, 1997), our findings raise the possibility that delta cells might play an important developmental role in controlling pancreatic expansion and maturation. Furthermore, we identified distinct endothelial cell states, including angiogenetic endothelial cells in neonatal pancreata that may reflect the increased supply of nutrients required by the rapidly replicating cells at this developmental stage; in particular, neonatal islets rapidly develop extensive glomerular-like circulatory structures which ensures the spatial proximity of endocrine cells to arterial blood (Bonner-Weir, 1988; Bonner-Weir and Orci, 1982; Cleaver and Dor, 2012). Here, application of modern, powerful computational tools also helped identify a previously uncharted landscape of cell-cell interactions. This included evidence of interactions between macrophages and endothelial cells in the neonatal pancreas, and possible ligand-receptor interactions involved in organ remodeling during growth (Geutskens et al., 2005). Our studies also revealed aspects of human endocrine pancreas development and regulation. The combination of adult and neonatal datasets allowed pseudotime analyses, and nominated candidate regulators and effectors of beta cell maturation, including age-dependent restriction of beta cell proliferation and development of secretory activities (Arda et al., 2016; Bonner-Weir, 2000). ISS analyses confirmed differences in islet size and intra-islet architecture between juvenile and adult pancreatic tissue, and provided evidence for possibly stereotyped dispersion of islets throughout the human pancreas, like in rodents (Pauerstein et al., 2017). In summary, our studies combine technical innovations to produce a human pancreas cell atlas that provides conceptual advances and reveals cellular, genetic, signaling and physiological mechanisms regulating pancreatic cells in health and disease.

## Supplementary Materials

**Supplementary Table 1.** Adult and juvenile human donor metadata.

| Sample ID | Sex | Age (years) | Pancreatic disease | Diabetes | Procurement lab | Pancreas location | # cells |
| --- | --- | --- | --- | --- | --- | --- | --- |
| AFHE365-body | F | 53 | None | None | Stanford, USA | Body | 22,288 |
| AFHE365-head | F | 53 | None | None | Stanford, USA | Head |  |
| AFHE365-tail | F | 53 | None | None | Stanford, USA | Tail |  |
| AFES448-head | M | 30 | None | None | Stanford, USA | Head | 34,167 |
| AFES448-midbody | M | 30 | None | None | Stanford, USA | Mid-Body |  |
| AFES448-body | M | 30 | None | None | Stanford, USA | Distal Body |  |
| AFES448-tail | M | 30 | None | None | Stanford, USA | Tail |  |
| AGBR024-body | M | 1.5 | None | None | Stanford, USA | Body | 33,196 |
| AGBR024-head | M | 1.5 | None | None | Stanford, USA | Head |  |
| AGBR024-tail | M | 1.5 | None | None | Stanford, USA | Tail |  |
| TUM-13 | F | 46 | Neuroendocrine Tumor | N/A | Munich, Germany | Tail | 5,233 |
| TUM-C1 | M | 77 | Pancreatic Ductal Adenocarcinoma | N/A | Munich, Germany | Body | 7,967 |
| TUM-25 | F | 59 | Mixed Muellerian Tumor | N/A | Munich, Germany | Body | 9,712 |

**Supplementary Table 2.**
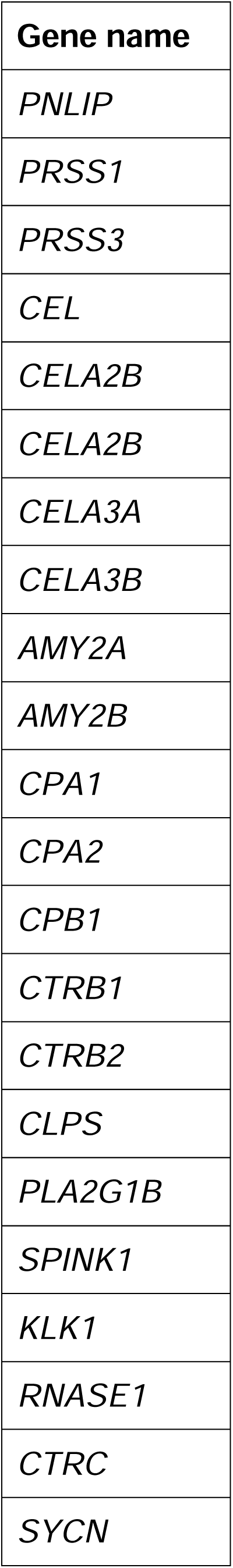
Table of digestive enzyme genes.

**Supplementary Table 3.** Neonatal human donor metadata.

| **Sample ID** | **Sex** | **Age**<br>**(days)** | **Pancreatic**<br>**disease** | **Diabetes** | **Procurement lab** | **Pancreas**<br>**location** | **# cells** |
| --- | --- | --- | --- | --- | --- | --- | --- |
| IIAM | M | 1 | None | None | Stanford, USA | N/A | 5,100 |
| ST19 | F | 1 | None | None | Stanford, USA | N/A | 5,428 |

**Supplementary Table 4.** ISS target gene list.

|  |  |  |  |
| --- | --- | --- | --- |
| ABCC8 | FABP5 | MEG3 | SERPINE1 |
| ACTA2 | FBLN1 | MUC5B | SLC4A4 |
| ADIRF | FLT1 | MUC6 | SLC8A1 |
| AMY2A | G6PC2 | PDK4 | SLCO2A1 |
| ANXA4 | GC | PIGR | SLPI |
| APOD | GCG | PIK3R3 | SOX10 |
| B2M | HLA-DRA | PKHD1 | SPARCL1 |
| BICC1 | IAPP | PLP1 | SST |
| CALD1 | ID1 | PLVAP | SYT8 |
| CD14 | INS | PMP22 | TENM2 |
| CD7 | INSR | PNLIP | TFF3 |
| CD74 | IRX2 | PNLIPRP1 | TGFBR3 |
| CDH19 | ITGA7 | PPY | THSD7A |
| CFTR | KCNMB2 | PRSS1 | TIMP3 |
| CHRM3 | KCNT2 | RBP4 | TRHDE |
| CPA2 | KRT19 | RBPJL | TRPM3 |
| CRYAB | LCN2 | REG3A | TTR |
| CTRB1 | LDB2 | REG3G | VWF |
| CUZD1 | LOXL4 | S100B | ZEB2 |
| DCN | LRIG1 | SCN7A | ZNF385D |
| FABP4 | MECOM | SCTR |  |

**Figure S1.**
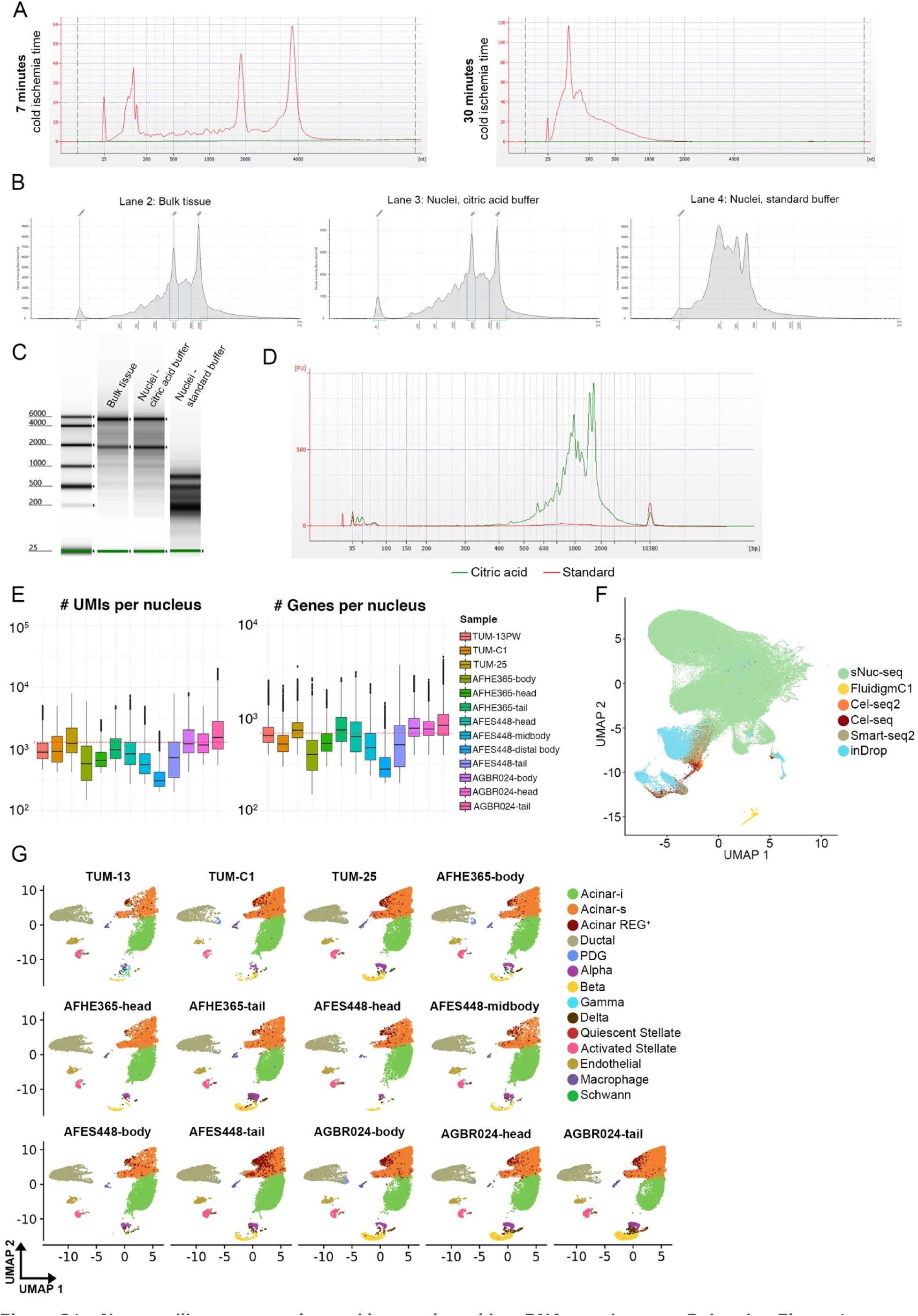
sNuc-seq library generation and integration with scRNA-seq datasets, Related to Figure 1. (A) Electropherogram of bulk RNA extracted from snap-frozen pig pancreatic tissue subject to either 7 or 30 minutes of total cold ischemia (B) Electropherograms of bulk RNA extracted from snap-frozen human pancreatic tissue (Bulk tissue). RNA was extracted from nuclei that were isolated from the same tissue as in lane 2 by using either a citric acid buffer or the standard buffer (lanes 3 and 4). (C) Gel view of the same samples as in (B). (D) Yield of cDNA from a sample processed with either the standard or the citric acid-based protocol. The same number of nuclei and PCR cycles were used for both conditions. (E) On the left, the boxplots show the distribution of Unique Molecular Identifiers (UMIs) per nucleus for each sample processed in this study. On the right, the boxplots show the distribution of genes per nucleus for each sample. The red dashed lines represent mean values (1,287 for UMIs and 692 for the genes). (F) Merged sNuc-Seq and previously published scRNA-seq datasets shown in a two-dimensional UMAP embedding before batch effect removal. (G) Following batch-effect removal, sNuc-seq data were split by sample of origin and shown in a two-dimensional UMAP embedding.

**Figure S2.**
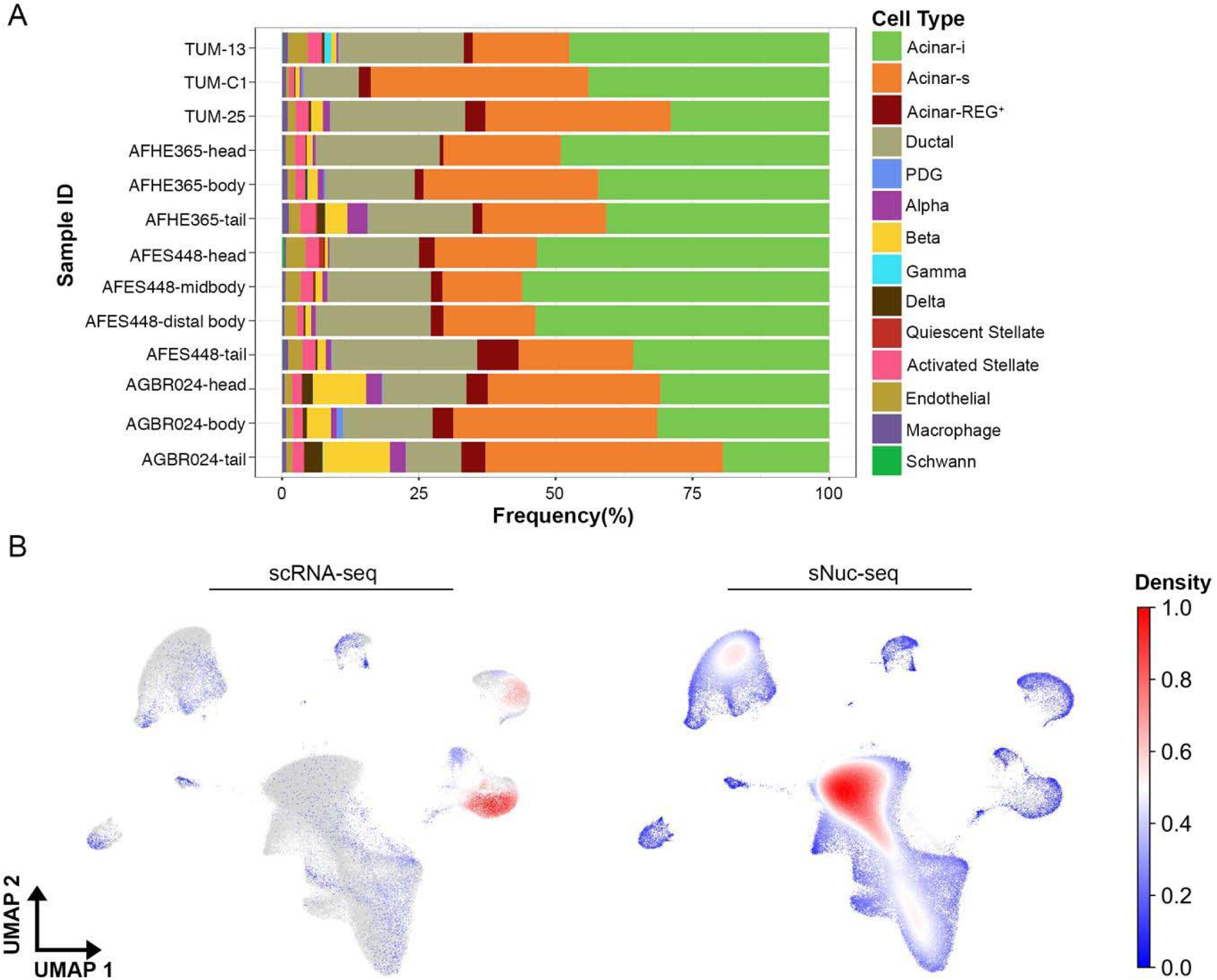
Different proportion of cells detected by sNuc-seq and scRNA-seq, Related to Figure 1. (A) Barplots showing the proportion of cell types identified in each sNuc-seq sample. (B) Gaussian kernel density estimation was used to calculate the density of cells and was represented in the UMAP embedding for the two distinct technologies, namely scRNA-seq and sNuc-seq. High density values indicate strong contribution of the cells to the overall dataset (i.e. exocrine cells have higher contribution in sNuc-seq and endocrine cells in scRNA-seq).

**Figure S3.**
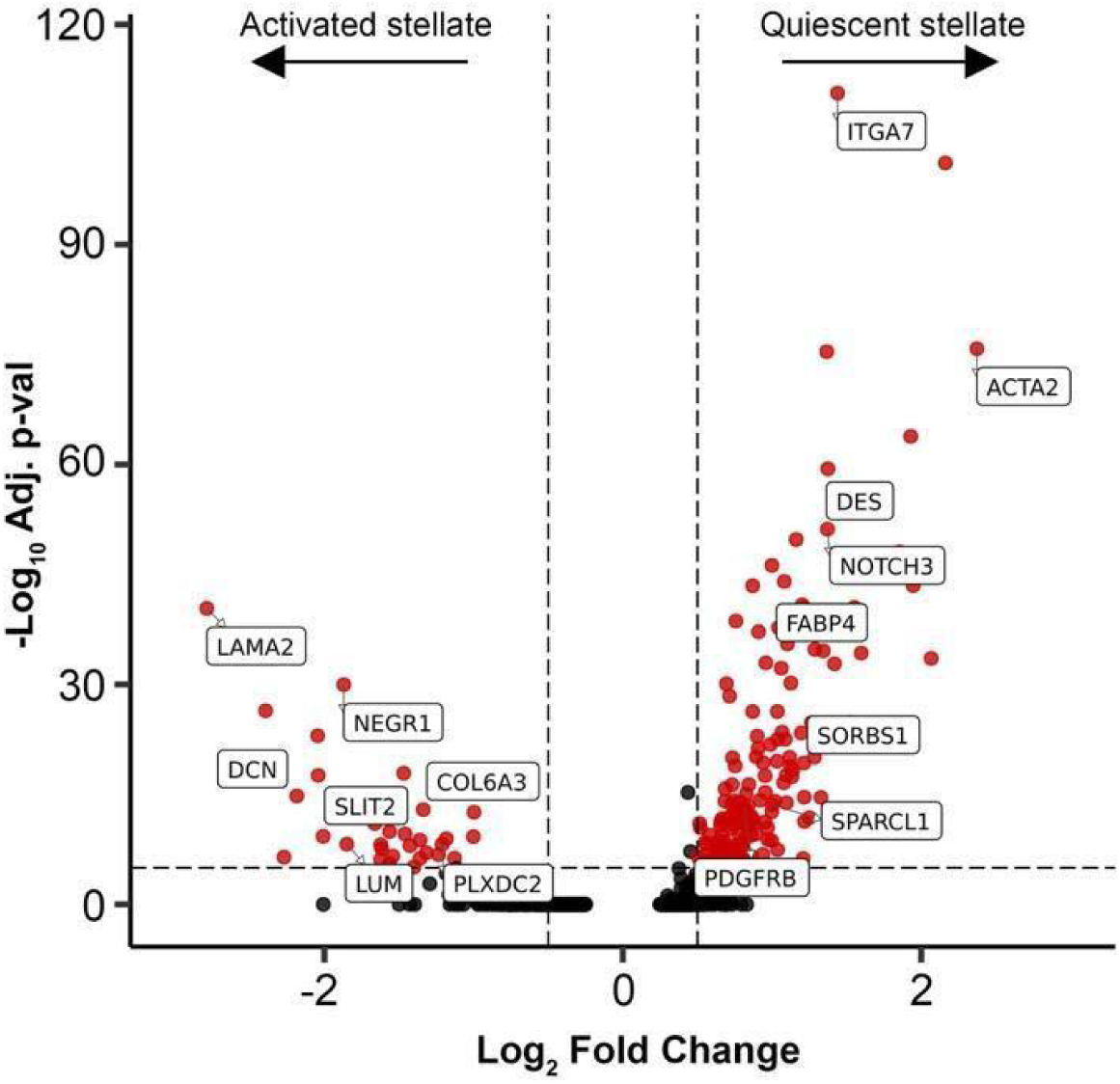
Differential gene expression between stellate cells, Related to Figure 3. Volcano plot showing differentially expressed genes between activated and quiescent stellate cells. Red dots represent genes with average log expression >0.5 and an adjusted p-value <0.05.

**Figure S4.**
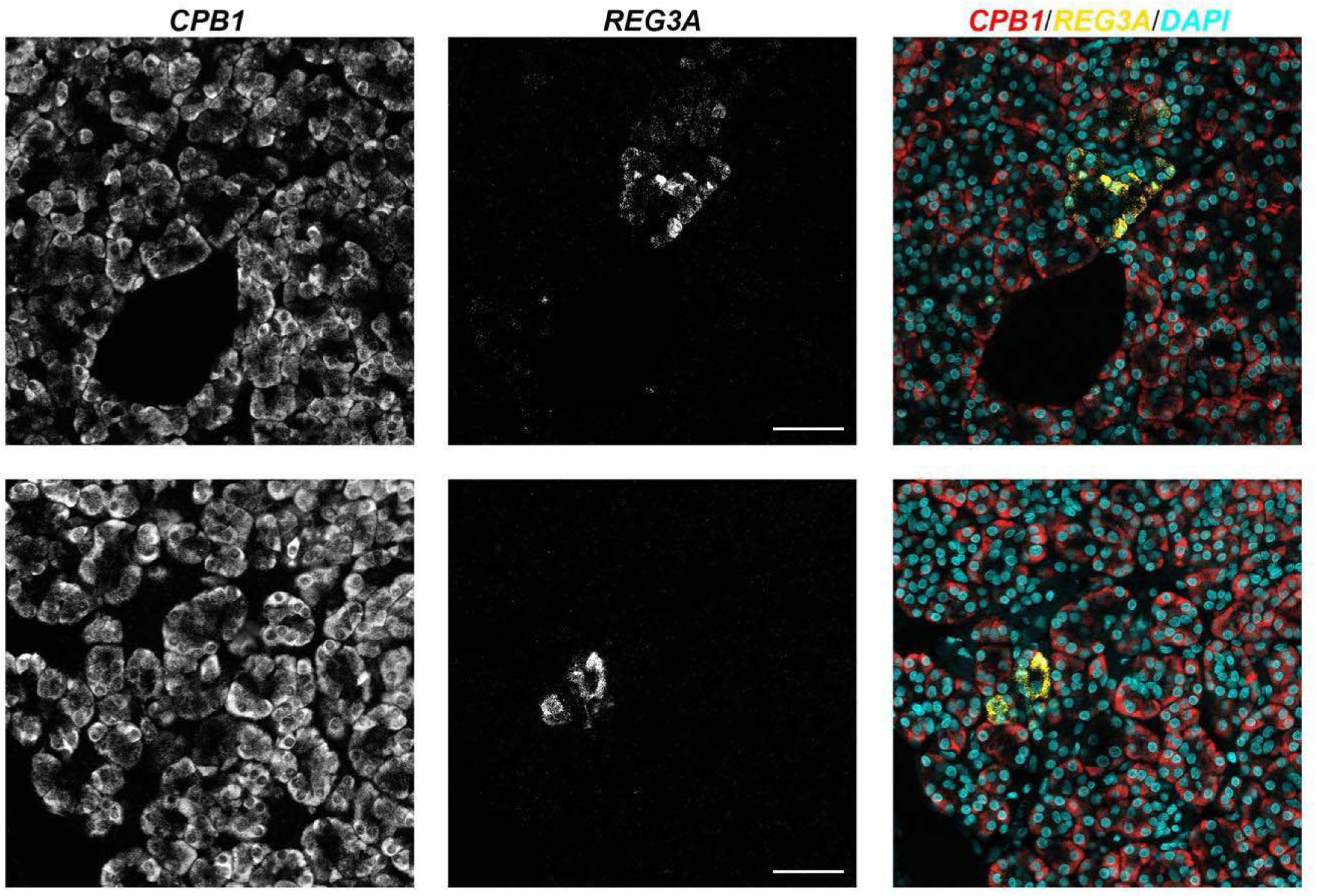
RNA-FISH in the healthy human pancreas, Related to Figure 4. Example image of RNA-FISH for *CPB1* and *REG3A* in the human adult healthy pancreas. Acinar-REG^+^ cells constitute a subset of CPB1^+^ acinar cells.

**Figure S5.**
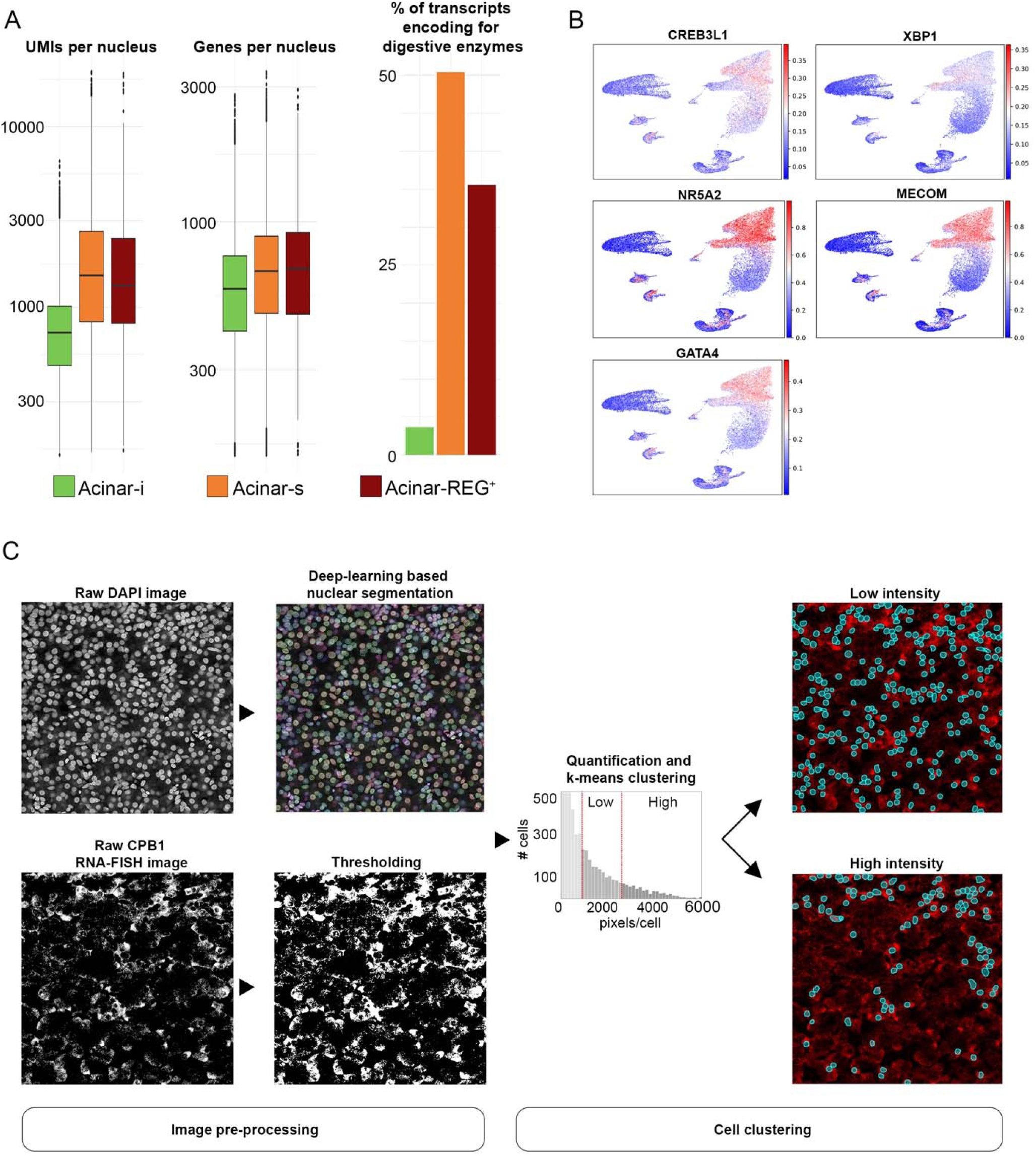
Characterization of acinar-i and acinar-s cells, Related to Figure 5. (A) Quantification of UMI per nucleus (left) and genes per nucleus (center) for the different acinar cell states. On the right, the percentage of transcriptome encoding for digestive enzymes (Table S2) is represented. (B) SCENIC regulons specifically active in the acinar-i and acinar-s cells. Above each plot, the transcription factor is indicated. (C) Pipeline applied for the quantification of differential RNA-FISH signal. Raw RNA-FISH images were thresholded to remove the background signal and a nuclear segmentation mask was generated by applying a deep-learning algorithm to the DAPI channel of the same image. Signal was quantified for each nucleus and the signal distribution was used to identify acinar cells expressing different level of digestive enzyme genes as explained in Materials and Methods.

**Figure S6.**
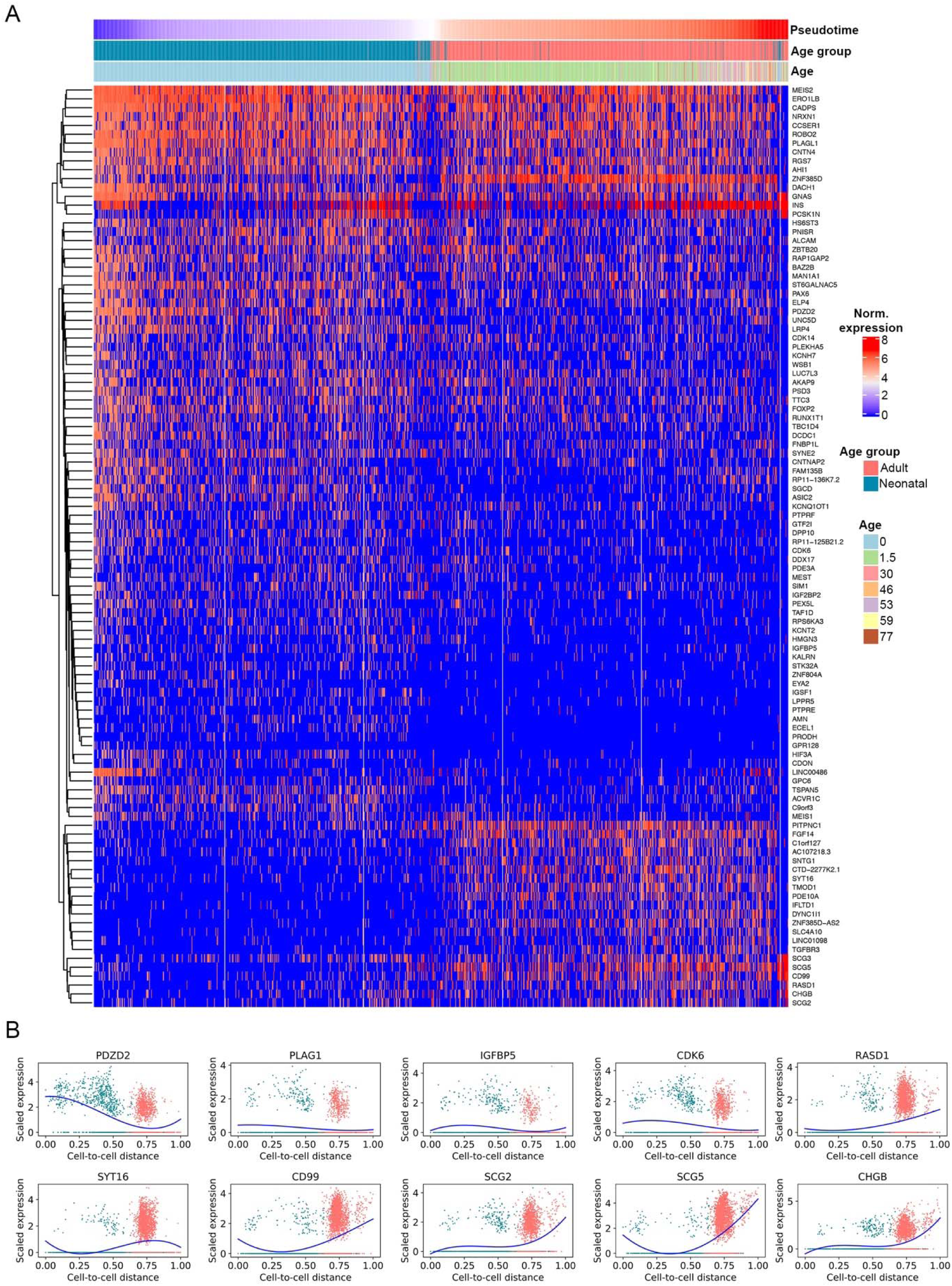
Gene expression changes in beta cell from neonatal to adult stage, Related to Figure 6. (A) Heat map showing gene expression changes across pseudotime reflecting beta cell maturation, from neonatal to adult. (B) The dots indicate the expression levels of individual cells colored by age type in the β-cell cluster. The blue lines approximate expression along the inferred trajectory by polynomial regression fits.

**Figure S7.**
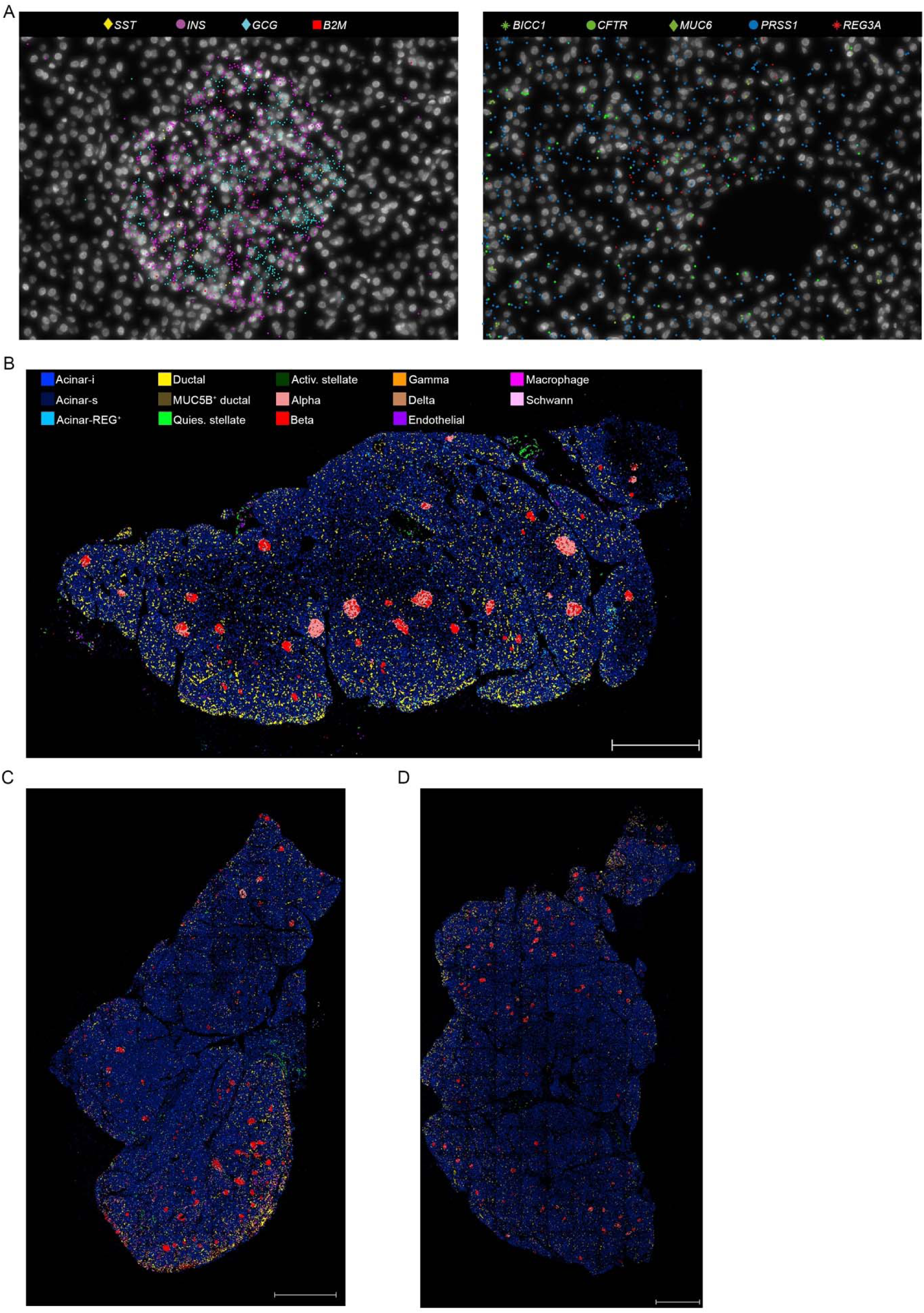
*In situ* sequencing signal and cell maps, Related to Figure 7. (A) On the left, localization of different marker genes in an endocrine islet (*SST* for delta cells, *INS* for beta cells, *GCG* for alpha cells, *B2M* for endothelial cells) as captured by ISS. On the right, markers for ductal (*BICC1, CFTR, MUC6*) and acinar cells (*PRSS1, REG3A*) as captured by ISS. (B) Cell map generated by SSAM from a tissue section of the sample AFES448-midbody. (C) Cell map generated by SSAM from a tissue section of the sample AGBR024-head. (D) Cell map generated by SSAM from a tissue section of the sample AGBR024-body. For the metadata, see Table S1. Scale bar = 1mm.

**Figure S8.**
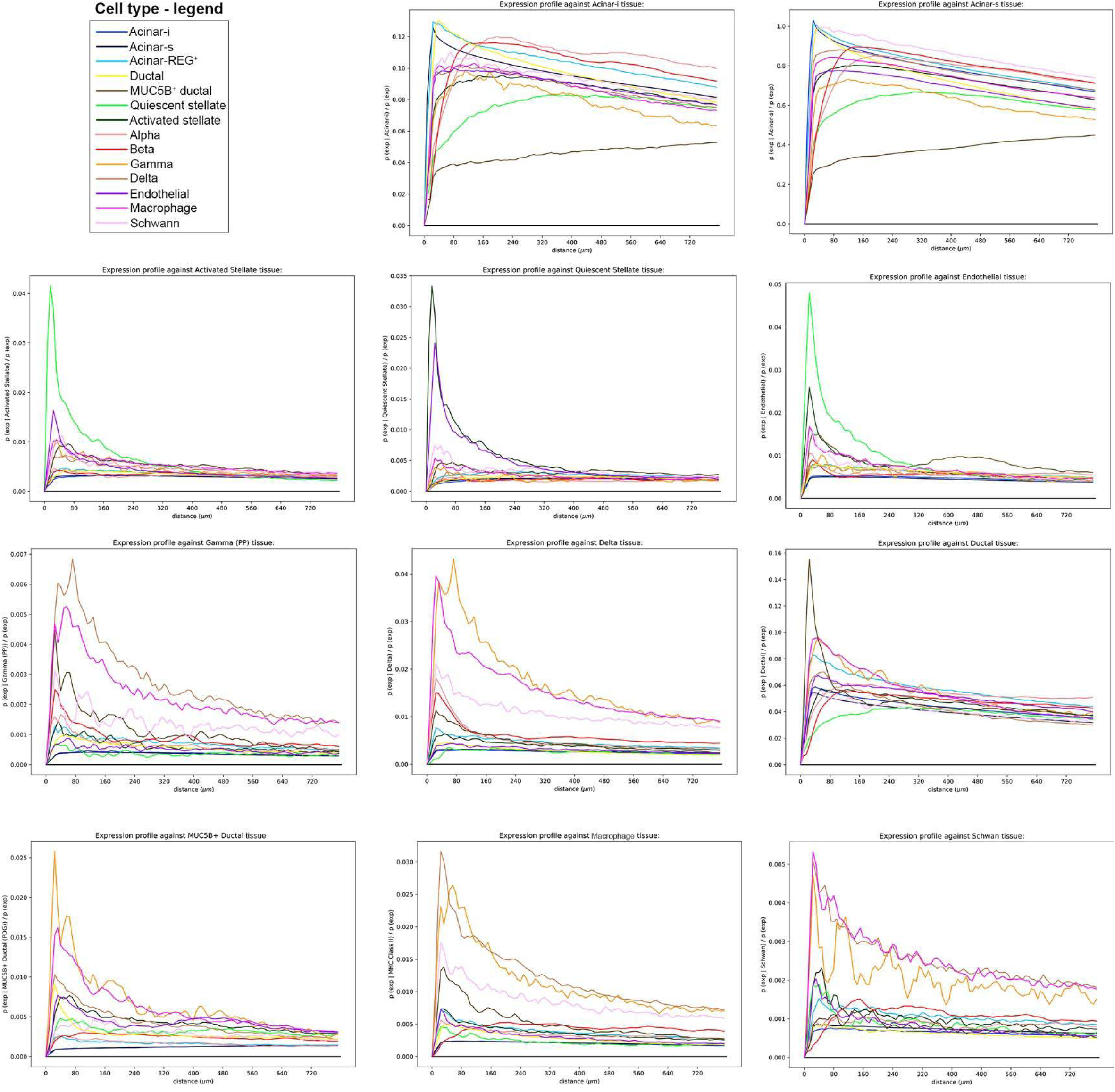
Statistical modelling of spatial relationship for different pancreatic cell types, Related to Figure 8. Line plot showing the results of the modelling analysis for 11 different cell types.

## Acknowledgements

The authors would like to thank organ donors and their families, David Ibberson (Heidelberg University), Ulrike Krüger (BIH, Berlin) and Marten Jager (BIH, Berlin) for NGS services, the Biomaterial Bank (MTBIO) of the Technical University Munich for support, Katharina Jechow (BIH, Berlin), Lorenz Chua (BIH, Berlin), Dr. Alison McGarvey (MDC, Berlin) for critically revising the manuscript, Jeongbin Park (BIH, Berlin) for his help with the analysis of ISS data and all the members of the Conrad lab for the constructive discussions. This study was funded by Human Cell Atlas (HCA) pilot studies of the Chan Zuckerberg initiative (Charité and Technische Universita□t Mu□nchen: CZI grant 2017-174170) and the European Marie-Sklodowska Curie Actions (EC no. 841755). The authors gratefully acknowledge the data storage service SDS@hd supported by the Ministry of Science, Research and the Arts Baden-Württemberg (MWK) and the German Research Foundation (DFG) through grant INST 35/1314-1 FUGG and INST 35/1503-1 FUGG. This publication is part of the Human Cell Atlas - www.humancellatlas.org/publications.

## Author contributions

C.C., R.E., S.K. and W.W. conceived the study. L.T. optimized the citric acid-based protocol, performed sNuc-seq, RNA-FISH and ISS experiments, analyzed the sNuc-seq data and interpreted the results. Y.H., R.B., K.S., S.B. procured human and pig samples. T.T. performed initial experiment in pig pancreas. A.A.K. and S.S. provided support and performed histology experiments. F.W.T. analyzed RNA-FISH images and S.L. provided support with data analysis and interpretation. O.D. performed trajectory analyses in adult and neonatal datasets and S.T. analyzed *in situ* sequencing data. R.E., C.C. and N.I. supervised experiment design and data interpretation. L.T., C.C., Y.H., S.K., R.E. wrote the manuscript with input from all the coauthors.

## Materials and Methods

### Human and pig pancreas samples

Samples from Stanford University were procured from non-diabetic cadaveric organ donors. All studies involving human specimens were conducted in accordance with Stanford University Institutional Review Board guidelines. Deidentified human pancreata were procured from previously healthy, non-diabetic donors with less than 12-hour cold ischemia time through the Center for Organ Recovery and Education, CORE, Pittsburgh, PA, USA), International Institute for the Advancement of Medicine (IIAM, Edison, New Jersey, USA), and National Diabetes Research Institute (NDRI, Philadelphia, PA, USA) as reported previously (Goodyer et al., 2012). Within minutes of removal from cold transportation media, tissue blocks of 2 cm × 1 cm × 0.2 cm were excised from 3-4 anatomic locations (i.e. head, body, mid-body, and tail) and then immediately transferred into liquid nitrogen to snap freeze. The frozen samples were shipped and stored at −80°C until they were used for nuclei isolation and sequencing. Samples from TUM were procured from non-diseased pancreatic tissue from patients undergoing partial pancreatectomy. Tissue blocks of 0.5 cm × 0.5 cm × 1 cm were collected immediately after removal of the pancreas, placed into cryo tubes and transferred into liquid nitrogen. The samples were stored at −196°C until they were used for sequencing. The study was approved by the hospital Ethics Committee (number 403/17S). To further dissect the preanalytical problems in procurement of pancreatic tissue for single-cell sequencing, we also sampled pancreatic tissue from healthy pigs sacrificed due to other reasons (approved by local authorities AZ .3-8-07, Regierung von Oberbayern, München, Germany) under completely standardized conditions. The pancreas was removed after the heartbeat had stopped and tissue blocks (0.5 cm × 0.5 cm × 1 cm) were sampled at different time points (15 min and 30 min cold ischemia time) and transferred into liquid nitrogen. The samples were stored according to the requirements of fixation solution/procedure. To check for morphological integrity of the tissue, a paraffin block and hematoxylin-eosin stained slide was produced from each sampling site and evaluated by two experienced pathologists (S.B. and K.S.).

### Nuclei isolation

Snap-frozen pancreatic tissue samples were cut into pieces <0.3 cm and homogenized with one stroke of “loose” pestle in 1 mL citric-acid based buffer (Sucrose 0.25 M, Citric Acid 25 mM, Hoechst 33342 1 μg/mL) using a glass dounce tissue grinder. The tissue was incubated on ice for 5 minutes and then homogenized with 5-10 more strokes. After further 5 minutes of incubation, tissue was homogenized with 3-5 strokes using the “loose” pestle and then 5 more strokes using the “tight” pestle. Homogenate was filtered through a 35-μm cell strainer and centrifuged for 5 minutes at 500 x g at 4°C. Supernatant was carefully removed, nuclei were resuspended in 1 mL of citric acid buffer and the centrifugation step was repeated. Nuclei were then resuspended in 300 μL of cold resuspension buffer (KCl 25 mM, MgCl_2_ 3 mM, Tris-buffer 50 mM, RNaseIn 0.4 U/μL, DTT 1mM, SuperaseIn 0.4 U/μL, Hoechst 33342 1 μg/mL). Nuclei were counted on a Countess II FL Automated Cell Counter, diluted to the desired concentration and immediately loaded on the 10X Chromium controller.

### 10X sample processing, library preparation and sequencing

Samples were prepared according to the 10x Genomics Single Cell 3′ v2 and 10x Genomics Single Cell 3′ v3 Reagent Kit user guide with small modifications. The nuclei were diluted using an appropriate volume of resuspension buffer without Hoechst (KCl 25 mM, MgCl_2_ 3 mM, Tris-buffer 50 mM, RNaseIn 0.4 U/μL, DTT 1mM, SuperaseIn 0.4 U/μL) for a target capture of 10,000 nuclei. After droplet generation, samples were transferred onto a pre-chilled 96-well plate (Eppendorf), heat-sealed and reverse transcription was performed using a Bio-Rad C1000 Thermal Cycler. After the reverse transcription, cDNA was recovered using the Recovery Agent followed by a Silane DynaBead clean-up step. Purified cDNA was amplified for 15 cycles before bead cleanup using SPRIselect beads (Beckman). Samples were quantified using an Invitrogen Qubit 4 Fluorometer. cDNA libraries were prepared according to the Single Cell 3′ Reagent Kits v2 and Single Cell 3′ Reagent Kits v3 guide with appropriate choice of PCR cycle number based on the calculated cDNA concentration. Final libraries were sequenced with the NextSeq 500 system in high-output mode (paired-end, 75 bp).

### Single-cell RNA sequencing data analysis

#### Alignment

Gene expression was quantified using the default 10X Cell Ranger v3 pipeline but we used a curated genome annotation of the GRCh37/hg19 Reference - 2.1.0 provided by 10X, in which the *INS-IGF2* gene sequence overlapping with the *INS* gene sequence was removed (Wernersson et al., 2015). Introns were annotated as “mRNA”, and intronic reads were included in expression quantification matrix.

#### Quality control and downstream analyses

During nuclei isolation, the cytoplasmic content of each cell (including mature mRNA) is released in the nuclei suspension. To reduce the background levels of this ambient RNA, nuclei were washed twice or three times before loading on the 10X Chromium controller. However, enzymatic digestive genes are highly expressed and known to be the source of contamination in bulk and scRNA-seq studies (Nieuwenhuis et al., 2019) hence we applied SoupX for background correction to the matrix generated by the 10X Cell Ranger v3 pipeline (Young and Behjati). In SoupX the ambient RNA expression is estimated from the empty droplets (i.e. droplets containing less that 10 UMI) and the expression of these genes is calculated and compared with their proportion in the ambient RNA profile. To calculate the contamination fraction, we used the *PRSS1* gene. The contamination fraction derived from the expression of *PRSS1* was used to calculate the fraction of each droplet expression corresponding to the actual cell. Finally, this fraction and the ambient profiles are subtracted from the real expression values to generate the background-removed expression matrices. Quality control (QC) analyses were performed on the basis of guidelines recently described (Amezquita et al., 2020). In particular, UMI and genes were filtered for each sample after visual inspection of QC metric diagnostic plots. In general, nuclei with a minimum number of genes between 150-400 and maximum number of genes between 2000-5000 were kept. Moreover, nuclei containing more than 3% of mitochondrial reads were excluded from downstream analyses. In addition to the general QC described above, we removed small clusters of nuclei co-expressing acinar and ductal markers, namely *CFTR* and *PRSS1*. Downstream analyses were performed using the R package Seurat version 3.0 and included also five previously published scRNA-seq datasets (Stuart et al., 2019). Each sNuc-seq dataset was scaled by library size and log-transformed (using a size factor of 10,000 molecules per cell). For each sample, the top 2,000 most variable genes were identified and the sNuc-seq and scRNA-seq datasets were integrated using the “FindIntegrationAnchors” and “IntegrateData” available in Seurat 3.0 (Stuart et al., 2019). Data were scaled to unit variance and zero mean and the dimensionality of the data was reduced by principal component analysis (PCA) (30 components) and visualized with UMAP (McInnes et al., 2018). Clustering was performed using the Louvain algorithm on the 30 principal components (resolution = 1.0). Small clusters including Schwann cells and MUC5B^+^ ductal cells were manually assigned. Cluster-specific markers were identified with the “FindAllMarkers” function and clusters were assigned to known cell types on the basis of their specific markers (described in the main text). Clusters that appeared to correspond to the same cell types were merged. The density map in Figure S4B was calculated and plotted using the “embedding_density” function of Scanpy version 1.4.2 (Wolf et al., 2018).

## Reconstruction of lineage relationships and trajectories in the adult pancreas dataset

To infer the lineage relationships and global transcriptomic similarity between different adult pancreatic cell types, we performed partition-based graph abstraction (PAGA) analysis that provides an interpretable graph-like map by measuring cluster connectivity (Wolf et al., 2019). PAGA was calculated using the tl.paga function implemented in Scanpy (v. 1.4.2) with an edge significance threshold of 0.6. The output of this function is an adjacency network where nodes are cell types and the edges represent connections between them. Here, the edge weights signify the confidence of a connection calculated based on a measure of cluster connectivity. Both PCA (tl.pca function implemented in Scanpy) and ICA (FastICA function implemented in sklearn.decomposition) (Pedregosa et al., 2011) were applied as linear dimension reduction methods leading to the same result. The pseudotemporal ordering of the cells was computed using the tl.dpt function of Scanpy by setting a root cell within the acinar-i cell population. Linear dimension reduction was performed using ICA (Pedregosa et al., 2011) and the calculated components were used to compute the neighbourhood of the single cell graph using the sc.pp.neighbors function using 50 nearest neighbors in an adaptive Gaussian kernel.

## Gene expression dynamics of beta cell maturation

The adult and neonatal datasets were merged using Seurat version 3.0 (Stuart et al., 2019) and the count matrices of beta cells were exported for further analyses in Scanpy (v. 1.4.2) (Wolf et al., 2018). Linear dimension reduction was performed using PCA followed by an unsupervised diffusion map analysis with the tl.diffmap() function on the first 15 neighbors. We removed acinar (Table S2) and ductal markers (*CFTR, BICC1, ANXA4*) as they represent contamination from other cell types and computed pseudotemporal ordering using the tl.dpt function of Scanpy by setting a root cell within the neonatal population (Wolf et al., 2018). A generalized additive model (GAM) was then used for modeling gene expression profiles as nonlinear functions of pseudotime for neonatal and adult lineages (Hastie, 2019). The top 105 genes that are a function of pseudotime are plotted as an annotated heatmap (Figure S11) (Gu et al., 2016).

### Regulon - SCENIC analysis

SCENIC (Aibar et al., 2017) is able to infer gene regulatory networks from single cell gene expression data through three main steps: (a) identification of co-expression modules between TF and putative targets; (b) within each co-expression module, derivation of direct TF-target gene interaction based on enrichment of TF motif in the promoter of target genes, as to generate “regulons”; (c) for each cell, the regulon activity score (RAS) is calculated. In this work, we applied the python implementation of SCENIC (pySCENIC) to downsampled datasets (30,000 nuclei) and the RAS was projected onto the UMAP embedding calculated by Seurat.

### Gene over-representation analysis

Symbol gene IDs were converted to Entrez gene IDs using the R package “annotables” (Turner). The Gene Ontology over-representation analysis was performed using the “enrichGO” function of the clusterProfiler R package (Yu et al., 2012) (using adjusted p-value <0.05 and average log(Fold Change) >0.25). The KEGG over-representation test was performed using the “enrichKEGG” function and the enrichment maps in Figure 2D and Figure S10A were generated using the “emaplot” function.

### Histology and RNA-FISH

To perform RNA-FISH in the human pancreas, we used thin snap-frozen (2 mm) biopsies for formalin fixation and paraffin embedding, reasoning that the fixation of the tissue would be faster due to the thinness of the tissue, limiting the degradation processes. Therefore, human pancreatic snap-frozen samples were fixed in 10% formalin at 4°C for 14-16 hours and paraffin-embedded. Sections (4 μm) were cut from FFPE pancreatic tissue and processed for RNA *in situ* detection using the RNAscope Multiplex Fluorescent Reagent Kit v2 according to the manufacturer’s instructions (Advanced Cell Diagnostics). RNAscope human probes used were: Hs-CPB1 (#569891-C3), Hs-RBPJL (#581131), Hs-AMY1A (#503551-C2, targeting also AMY1B, AMY1C, AMY2A and AMY2B), Hs-REG3A (#312061). RNA-FISH images were acquired on a Leica SP8 confocal laser-scanning microscope equipped with a 40x/1.30 oil objective (Leica HC APO CS2).

### RNA-FISH image analysis

Automated nuclei instance detection and segmentation were implemented and performed using a deep learning object detection and instance segmentation workflow based on the Mask R-CNN architecture (He et al., 2017). The neural network was initialized using pre-trained models trained on the Microsoft COCO: Common Objects in Context dataset (Lin et al., 2014) and fine-tuned on curated datasets of nuclei images. Nuclei images on the DAPI channel were used as inputs for the neural network to produce segmentation for each individual nucleus. The nuclei sizes were calculated using these segmented nuclei masks, and objects <150 pixels were filtered out and excluded from subsequent analyses.

To perform transcript abundance analysis, the RNA-FISH channels were thresholded and binarized by computing the gray-level moments of the input images as implemented in Fiji. Transcript abundance was estimated by overlaying the nuclei masks on the thresholded probe channels and calculating the number of pixels within each mask. In order to account for transcript signals that are predominantly localized outside of the nuclei masks, we expanded the nuclei masks by morphological dilation (3 iterations using a 7×7 elliptical kernel) as implemented in OpenCV (Bradski, 2000) prior to quantification. We then performed k-means clustering on the frequency distributions of pixel counts per cell (nucleus) to identify and separate the cells into population classes (e.g. high, low, and negative expression/abundance). A cluster number of 3 was selected for the FISH signals to better capture gradual differences between cells.

### In situ Sequencing by CARTANA

Sections (4 μm) were cut from FFPE pancreatic tissue prepared as described for RNA-FISH. Four DNA probes for each target gene (6 base sequences with a minimum of 2 bases difference between all barcodes) were designed and supplied by CARTANA. Tissue fixation, reverse transcription, probe ligation, rolling circle amplification and fluorescence labeling were performed according to the manufacturer instructions (Neurokit 1010-01, CARTANA, Sweden). To reduce lipofuscin autofluorescence, 1X Lipofuscin Autofluorescence Quencher (Promocell) was applied for 30 seconds before fluorescence labeling. Samples were then shipped to CARTANA (Solna, Sweden) for the sequencing step. The result table of the spatial coordinates of each molecule for the 83 targets together with the reference DAPI image per sample were provided by CARTANA.

### SSAM and neighborhood analysis

For the analysis of the ISS data, the topographies of the primary tissue samples were reconstructed using the SSAM tool. In the first step, a list of cell-type-wise mRNA expression signatures was compiled based on prior knowledge and by-eye evaluation of the ISS data. To remove apparent noise from the dataset, all mRNA spots with a critical distance above 10um to its same cell type nearest neighbor were discarded (amounting to exclusion rates of 51%, 65%, 50%, and 64% of the respective samples). In the next step we created 83 gene-wise integrated mRNA expression densities using SSAM’s kernel density estimation (KDE) algorithm with a Gaussian kernel and a bandwidth parameter of 3um. To infer the spatial tissue composition, these spatial expression maps were integrated further using SSAM’s cell type mapping function on the pre-compiled cell-type/mRNA expression matrix. Each pixel was assigned the cell type that maximized the Pearson correlation between the expected expression profile and the inferred local expression mRNA densities. A pixel-wise assignment threshold of total expression of 0.005 was applied to discard low-confidence regions. Areas with a correlation value below 0.3 were discarded as inconclusive.

For the neighborhood analysis, a circular area with a radius of 800um was considered around each pixel. This area was subdivided into 200 equal-spaced ring-shaped distance intervals. The average surface area of the different tissue types inside each distance interval could then be correlated to the tissue type present at the center pixel. This resulted in an exhaustive, pixel-wise co-occurence map per combination of tissue type and per distance interval. The mean values across the spatial dimension of all map pixels covered by a certain cell type was then used to create a probability matrix of co-occurence of all tissue types and at all different distance intervals. The distance-occurence profile was plotted for all cell-cell combinations to recover recurring spatial patterns in cellular topography. For each visualization plot, a vortex tissue needed to be chosen as a central anchor point to plot the other tissues against. Then, a co-occurrence profile between the vortex tissue and the peripheral tissue was created by calculating the distance-wise ratios between the occurrence probability of the peripheral tissue around the vortex tissue and the expected global occurrence probability of the peripheral tissue. When plotted against the distance measure, this profile can be interpreted as the factor of increase or decrease of observing a given tissue type after learning the tissue type of your current location.

